# Large-scale field phenotyping using backpack LiDAR and GUI-based CropQuant-3D to measure structural responses to different nitrogen treatments in wheat

**DOI:** 10.1101/2021.05.19.444842

**Authors:** Yulei Zhu, Gang Sun, Guohui Ding, Jie Zhou, Mingxing Wen, Shichao Jin, Qiang Zhao, Joshua Colmer, Yanfeng Ding, Eric S. Ober, Ji Zhou

## Abstract

Plant phenomics is widely recognised as a key area to bridge the gap between traits of agricultural importance and genomic information. A wide range of field-based phenotyping solutions have been developed, from aerial-based to ground-based fixed gantry platforms and handheld devices. Nevertheless, several disadvantages of these current systems have been identified by the research community concerning mobility, affordability, throughput, accuracy, scalability, as well as the ability to analyse big data collected. Here, we present a novel phenotyping solution that combines a commercial backpack LiDAR device and our graphical user interface (GUI) based software called CropQuant-3D, which has been applied to phenotyping of wheat and associated 3D trait analysis. To our knowledge, this is the first use of backpack LiDAR for field-based plant research, which can acquire millions of 3D points to represent spatial features of crops. A key feature of the innovation is the GUI software that can extract plot-based traits from large, complex point clouds with limited computing time and power. We describe how we combined backpack LiDAR and CropQuant-3D to accurately quantify crop height and complex 3D traits such as variation in canopy structure, which was not possible to measure through other approaches. Also, we demonstrate the methodological advance and biological relevance of our work in a case study that examines the response of wheat varieties to three different levels of nitrogen fertilisation in field experiments. The results indicate that the combined solution can differentiate significant genotype and treatment effects on key morphological traits, with strong correlations with conventional manual measurements. Hence, we believe that the combined solution presented here could consistently quantify key traits at a larger scale and more quickly than heretofore possible, indicating the system could be used as a reliable research tool in large-scale and multi-location field phenotyping for crop research and breeding activities. We exhibit the system’s capability in addressing challenges in mobility, throughput, and scalability, contributing to the resolution of the phenotyping bottleneck. Furthermore, with the fast maturity of LiDAR technologies, technical advances in image analysis, and open software solutions, it is likely that the solution presented here has the potential for further development in accuracy and affordability, helping us fully exploit available genomic resources.

## Introduction

With the rising world population, crop production needs to double by 2050 (UN Food & Agriculture Organization, 2009). To address this growing challenge of global food security, it is important to identify plants with desired traits to improve yield, resource use efficiency, quality, stress resistance and adaptation, and with a smaller environmental footprint (Powlson et al., 2014; Zhang et al., 2018; Swarbreck et al., 2019). Furthermore, the stability of the selected traits must be verified in the field over multiple seasons and locations (Sadras and Slafer, 2012; Griffiths et al., 2015; Reynolds and Langridge, 2016). For example, quantitative measurements of yield-related traits such as plant height, growth rate, canopy coverage and spikes per unit area can be used to indicate and explain variations in yield stability in different environments (Sadras and Richards, 2014; AHDB, 2015; Valluru et al., 2017; Furbank et al., 2019). The cost of genotyping has decreased dramatically in recent years, allowing genetic analysis of large populations (Cobb et al., 2013; Crain et al., 2016). However, field phenotyping on a large scale under realistic field conditions remains the bottleneck in genotype-phenotype association studies for crop improvement (Furbank and Tester, 2011; Zhao et al., 2019; Yang et al., 2020). Both large-scale data acquisition and phenotypic analysis of multiple traits at different time points and trial locations are still challenging, but often it is the meaningful phenotypic information most needed by breeders and crop researchers (Fiorani and Schurr, 2013; Tardieu et al., 2017; Furbank et al., 2019).

To relieve this current bottleneck and address the above challenges in field phenotyping, much attention has been placed upon the applications of remote sensing, internet of things (IoT), robotics, computer vision, and machine learning, resulting in a rapid technical progress in recent years (Pieruschka and Schurr, 2019; Zhao et al., 2019; Yang et al., 2020). A range of solutions have been developed, including the use of unmanned aerial vehicles (UAVs) and manned light aircraft for studying performance-related traits across fields (Bauer et al., 2019; Holman et al., 2019; Harkel et al., 2020); stationary gantry systems for deep phenotyping in fixed areas (Vadez et al., 2015; Kirchgessner et al., 2017; Virlet et al., 2017; Burnette et al., 2018); ground-based vehicles equipped with integrated sensor arrays to study canopy-related traits (Deery et al., 2014; Barker et al., 2016; Jimenez-Berni et al., 2018); hand-held or distributed sensing devices to measure various phenotypes during key growth stages (Hirafuji and Yoichi, 2011; Crain et al., 2016; Zhou et al., 2017c; Reynolds et al., 2019a). These methods possess diverse advantages and disadvantages concerning throughput, accuracy, mobility, affordability, scalability and, more importantly, biological relevance (Fritsche-Neto and Borém, 2015; Furbank et al., 2019; Pieruschka and Schurr, 2019; Reynolds et al., 2019b; Roitsch et al., 2019). To choose a field phenotyping approach is naturally depending on the nature of the research question; but despite the rapid methodological progress, gaps in large-scale field solutions remain.

Among recent field-based methods, Light Detection and Ranging (LiDAR) has attracted much attention as it provides information on plant morphological and structural features that are difficult or costly to quantify through traditional approaches (Lin, 2015; Stovall et al., 2017). As an active remote sensing technique, LiDAR computes the distance from laser scanners to a given target using pulsed laser beams, through which three-dimensional (3D) geometric features of the target can be recorded in point cloud datasets (Arnó et al., 2013). LiDAR-based tools have been successful in overcoming issues related to natural illumination and occlusion, which have been problematic for many field-based measurements (Sun et al., 2018; Jin et al., 2019). Although point clouds produced by LiDAR can be subject to noise and imbalanced densities (Bucksch et al., 2009), newly developed open-source analysis libraries such as WhiteboxTools (Lindsay, 2016) and Open3D (Zhou et al., 2018) can be utilised to conduct 3D points processing. However, these libraries were developed for generic 3D analysis, which requires experienced developers with computer vision and biological backgrounds to develop tailored solution to analyse specific LiDAR data, limiting their use by the plant research community.

LiDAR devices can be roughly classified into three types: airborne, fixed terrestrial and mobile (Hosoi and Omasa, 2009; Lin, 2015). Plant characters that have been estimated include: crop height, biomass, and canopy structure (Omasa et al., 2007; Naito et al., 2017; Harkel et al., 2020); leaf number, shape, and the plant capacity to intercept solar radiation (Sun et al., 2018; Jin et al., 2019); and grain yield (Jimenez-Berni et al., 2018; Li et al., 2020b). LiDAR-generated point clouds have also been used to improve parameterisation of crop models, enabling *in silico* testing to optimise trait combinations in breeding and crop growth simulation (Reynolds and Langridge, 2016; Wang et al., 2017; Walter et al., 2019). In comparison with alternative approaches that can record 3D plant traits such as Structure from Motion (SfM) (Duan et al., 2016), time-of-flight (Paulus, 2019), micro-computed tomography (Wu et al., 2019), and photogrammetry techniques (An et al., 2016; Holman et al., 2016), for high-throughput field studies, LiDAR provides a more reliable solution in scalability, accuracy, and relatively low sensitivity to changeable light conditions in the field.

Despite these advantages, there are several problems associated with current LiDAR techniques in field phenotyping. Airborne LiDAR (Li et al., 2015; Harkel et al., 2020) typically requires larger multi-rotor UAVs with sufficient payload capacity (normally >5 kg), which requires special trained pilot and local aviation authority’s clearance, adding to hardware and operating costs. Also, big drones generate strong downdraft that disrupts canopies when flying them at low altitudes to acquire high-resolution imagery. Fixed terrestrial LiDAR (Omasa et al., 2007; Stovall et al., 2017; Guo et al., 2018), on the other hand, is placed closer to plants and can generate high-resolution models. Nevertheless, this type of system requires more time to set up to capture spatial features, limiting its applications in large-scale phenotyping. Mobile LiDAR (Arnó et al., 2013; Araus and Cairns, 2014; Deery et al., 2014; Jimenez-Berni et al., 2018; Deery et al., 2020) includes handheld, backpack, and devices mounted on specialised phenotyping vehicles (e.g. Phenomobile), which can cover large trial areas. The main drawbacks of vehicle-mounted LiDAR are the costs of purchasing hardware, operating and maintenance, as well as the ability to access agricultural fields with difficult conditions or rugged terrain. Handheld LiDAR devices are lightweight and easy to use, but usually are equipped with low-cost laser sensors, limiting their capability to acquire high-quality and large-scale models (Hyyppä et al., 2020; Jin et al., 2021).

The backpack LiDAR (Masiero et al., 2018; Hyyppä et al., 2020; Su et al., 2020) has been applied successfully to forestry studies and land surveillance in recent years, showing promise for field-based crop research. Compare with other LiDAR systems it has good mobility relatively lightweight (normally around 10 kg), and highly integrated in hardware, which means that it is easy to operate and maintain. Because the laser scanner can be used in close proximity to plants (< 3 m), it can generate high-quality 3D models with up to 10 mm precision, depending on the laser sensor and the selected mapping mode. Depending on the laser scanner equipped, backpack LiDAR system could have an effective scan range of over 200 m, useful for phenotyping in forestry or orchard plantations, as well as large experimental areas for crop plants. Backpack LiDAR also provides an accurate spatial positioning system (i.e. a global navigation satellite system, GNSS), customised for field mapping at walking speed to enable an accurate 3D reconstruction (Masiero et al., 2018). As LiDAR technology has been maturing rapidly in recent years, it is likely that costs will decrease and this type of equipment could become more accessible for the research community (Guo et al., 2018; Panjvani et al., 2019; Jin et al., 2021).

Notably, the analytic software for LiDAR-based technologies is as important as the hardware. One limitation of many LiDAR systems is the lack of widely available, open analytical software solutions that can extract biologically relevant information from the large point cloud data (Lin, 2015; Zhao et al., 2019; Yang et al., 2020), preventing non-expert users from taking advantage of this technology for rapidly modelling crop structural features and mining phenotypic information to study spatial and temporal changes (Ubbens et al., 2018; Panjvani et al., 2019; Ward et al., 2019).

Here, we introduce an integrated solution that combines a backpack LiDAR system with a dedicated open-source analytic software called CropQuant-3D, developed for processing 3D point cloud datasets collected by the backpack LiDAR. The system combines 2D/3D image analysis with Discrete Fourier Transform to derive plot-based measurements of key structural traits such as crop height, canopy structure, and biomass. To our knowledge, this is the first use of backpack LiDAR for field-based plant phenotyping. We therefore developed a range of technical applications to utilise the device in the field, including the mapping protocol for cereal crops and the quick assessment of the data quality at different sites. In a case study of wheat (*Triticum aestivum* L.), we describe the application of the backpack LiDAR and CropQuant-3D to quantify varietal responses to three levels of nitrogen (N) fertilisation of eleven Chinese winter wheat varieties from the ‘Zhenmai’ and ‘Ningmai’ populations. By combining 3D trait analysis and yield data, we produced a performance matrix to rank and evaluate genotypic differences in N responses, resulting in the classification of four N response types. In particular, to ensure that our work could reach the broader research community, we have expanded CropQuant-3D to analyse point clouds generated from other sources such as gantry-mounted LiDAR and unmanned aerial vehicles (UAVs). Furthermore, a graphical user interface (GUI) of CropQuant-3D was developed to package all the analysis algorithms, so that users without any computing background could utilise the system in their research. The software, source code organised into modules, and testing data are freely accessible via our GitHub repository for the research community.

## Results

### In-field mapping protocol using the backpack LiDAR

Because limited research has been conducted on the use of backpack LiDAR in phenotyping, we therefore developed a range of technical applications to utilise the device in the field, including the optimal distance to map cereal crops, the design of mapping routes and angles, the quick assessment of the data quality, and the calibration method for 3D mapping at different sites. For example, a grid-style mapping approach was designed to routinely phenotype the large field trial in this study (red arrows in **Fig. 1a)**. We first recorded the 3D geo-coordinates of the trial area using a real-time kinematic (RTK) base station, which logged satellite-based positions with ± 5 mm error range in 3D (**Fig. 1b**). Then, a LiDAR operator walked around the perimeter of each N treatment block in the experimental field to image the entire experiment from different angles. Due to the scan range of the LiDAR, we did not need to walk around each individual plot in the N treatment block, saving significant time in operation. On average, it took the LiDAR operator 20-25 minutes to map an experiment field of 0.5-ha, equivalent to a mapping speed of around 1.2 ha per hour. To study canopy structural responses to different N treatments, we focused on the growth stages between heading (GS51-59) and grain filling (GS71-89) when crop canopy was largely established (Zadocks et al., 1974).

**Figure 1:**
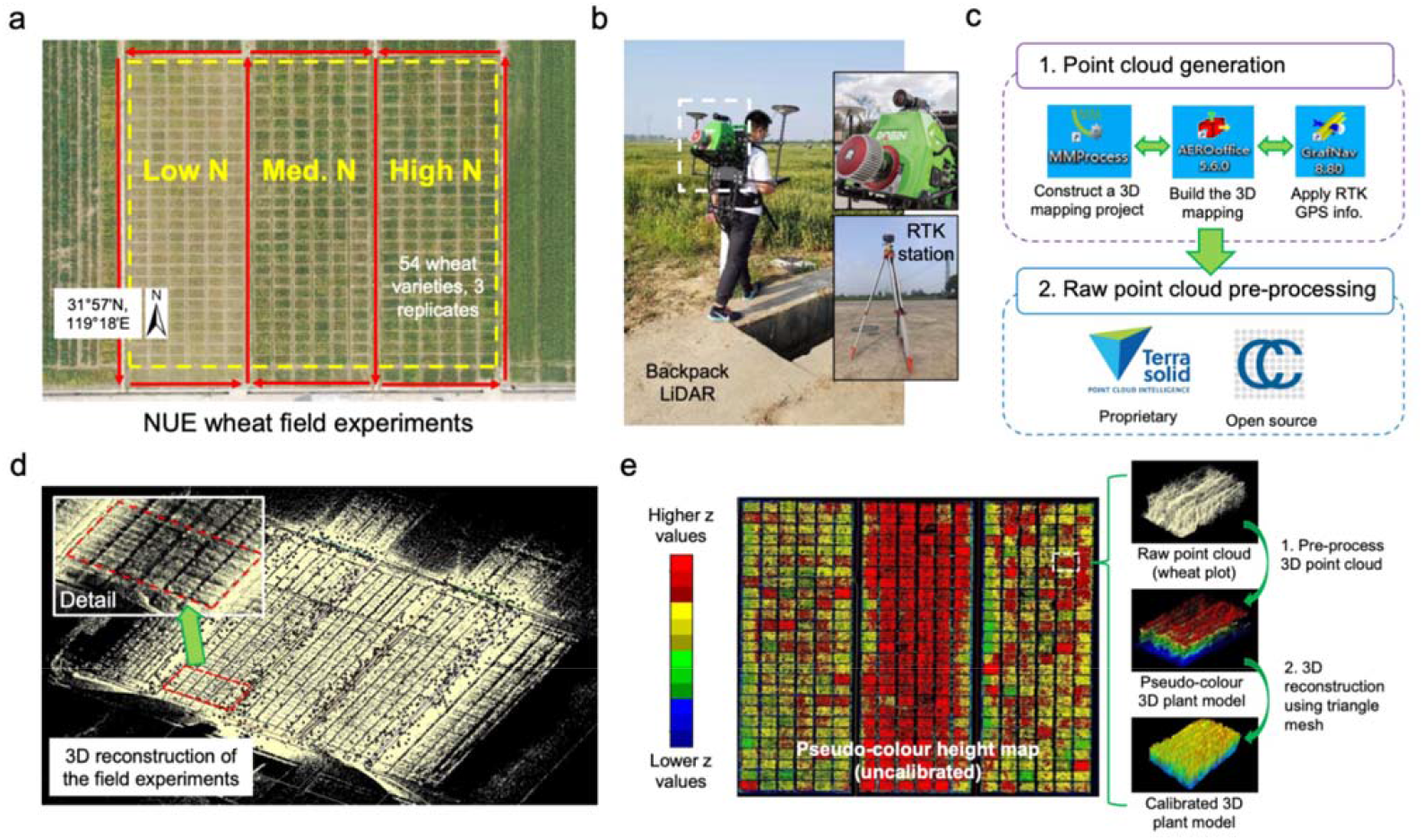
The data acquisition procedure using a backpack LiDAR device together with raw point cloud data generated through pre-processing a LiDAR-acquired 3D point cloud file. (**a**) An overhead orthomosaic image of the field trial area showing 486 six-metre winter wheat varieties with three levels of N fertilisation treatments (i.e. 0, 180, and 270 kg N ha-1). Red arrows represent the grid-style mapping method carried out by a LiDAR operator outside the plots. (**b**) The backpack LiDAR device (ROBIN Precision) and a real-time kinematic (RTK) base station used for 3D field phenotyping. (**c**) A high-level workflow of the pre-processing software used to generate GPS-tagged point cloud data collected by the backpack LiDAR. (**d**) The raw point clouds generated for the trial area. (**e**) Initial height-based analysis with uncalibrated 3D points, which were coloured according to GPS-tagged z values, and example plot-level images using raw 3D points, height values, and triangle mesh.

### Data pre-processing to generate 3D point clouds

According to standard practice in processing 3D points (Kachamba et al., 2016; Duan et al., 2017; Sun et al., 2018), we used the bundled pre-processing software to generate GPS-tagged 3D point clouds collected by the LiDAR (**Fig. 1c**). The bundled software we used are: MMProcess to build up a 3D mapping project, AERO-office to define the mapping path, and GrafNav to associate RTK GPS signals with the path. To select, visualise, and export point clouds of the fields, we chose to use the open-source CloudCompare software (Girardeau-Montaut, 2015). The same tasks can also be accomplished by using proprietary TerraSolid software (Korzeniowska and Łącka, 2011).

Because the backpack LiDAR device we used has an effective scan range of around 200 m (over 180 million points were collected in a single field mapping), the mapped area (over 1.5 ha, **Fig. 1d**) was much larger than the experiment region (i.e. the combined area of the 486 wheat plots, 0.5 ha). Hence, we used RTK-recorded geo-coordinates to delineate regions of interest (ROI) and facilitate our routine processing. After defining the ROI for the experimental region (over 45 million points retained for 486 plots, around 90,000 points per plot), all 3D points were visualised and coloured according to their z values (**Fig. 1e**). Terrain adjustment (e.g. slope removing and ground-level filtering) is a standard process for height estimates from elevation models in large-scale land surveillance and forestry research, as slopes and terrain features can lead to problematical height mapping. However, there are no standardised approaches designed for such adjustment in relatively small-scale crop fields; hence, we have incorporated a customised solution. Still, a preview of data prior to terrain adjustment enabled us to: (1) pseudo-colour raw point clouds for quick assessment; (2) perform initial comparisons of experiments at multiple sites; and (3) select ROI to facilitate plot-level 3D sampling.

### A comprehensive pipeline for traits analysis

To carry out routine 3D point processing and trait analysis using LiDAR-collected point clouds, we developed a comprehensive analysis pipeline and a GUI-based software application. **Figure 2** shows a high-level workflow of the analysis pipeline, which consisted of six steps: data selection, normalisation, the generation of crop canopy height model (CHM), plot segmentation, 3D trait analysis, and export of the analysis results:

**Figure 2:**
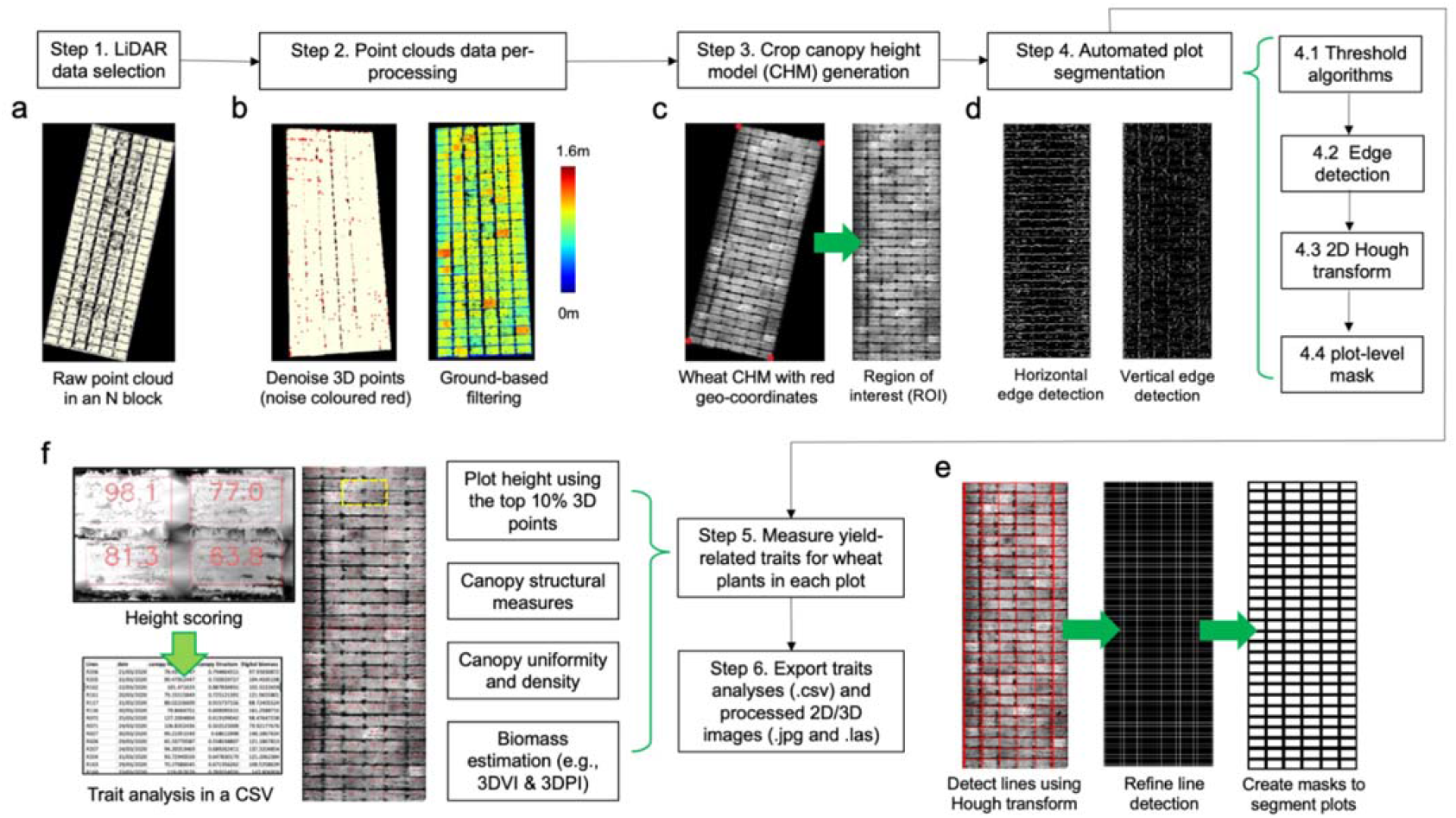
A comprehensive analysis pipeline established for processing LiDAR-acquired point clouds and measuring yield-related traits in 3D. (**a**) Select a pre-processed point cloud file (in LAS format). (**b**) Remove outliers (coloured red) in the point cloud, followed by filtering methods to differentiate ground-based terrain (e.g. soil level below the crop) and above-ground (crops) 3D points. (**c**) Generate a 2D canopy height model (CHM) and define the region of interest (ROI, denoted by the four red markers) using geo-coordinates collected by the ground based RTK station. (**d & e**) Detect horizontal and vertical edges using the Sobel operator, followed by the application of 2D Hough transform to produce a binary mask to segment plots in the field experiments. (**f**) Measure and export 3D trait analysis results for each plot, including measured traits (CSV), processed images (JPG), and processed point cloud (LAS).

1. *Steps 1*: a pre-processed point cloud file (in LAS format) was selected (**Fig. 2a**). Because LiDAR-collected point clouds is likely to be noisy and uncalibrated (with slopes and terrain features of the experimental field), we developed a process to normalise the 3D points (*Steps 2&3*). To remove noise, we followed a published method (Su et al., 2019), which calculates the average distance between a given 3D point and its neighbouring points (*avg*.). If the distance (*k*) between the point and its neighbouring 3D points (defaulted to 50) is greater than *avg*. + *k* × *std*. (where *std*. is one standard deviation of the mean of all the distances), the point will be classified as an outlier. In our case, all identified outliers were coloured red and removed from the following analysis (**Fig. 2b**).
2. *Step 2*: after denoising, a filtering method was applied to separate ground-level and above-ground (e.g. crops) 3D points by applying the `LidarGroundPointFilter` function in WhiteboxTools (Lindsay, 2016), including (1) ground-based slope normalisation; (2) a subsequent k-nearest neighbours, kNN, (Lowe, 2004) to identify neighbouring points within a defined *radius* (defaulted to 2) to examine height differences; and (3) a classification method to classify ground-level points and above-ground points. The use of the function resulted in a flattened ground plane of the field experiment, enabling precise measurements of above-ground 3D points, i.e. crop plants. The output of *Step 2* is saved in a new LAS file with all the ground-level points assigned with *zero* z-values (dark blue) and above-ground points assigned with height values in centimetre (cm).
3. *Step 3*: a key step in the pipeline used to generate a CHM for automated 3D trait analysis. First, because the density of LiDAR-collected point clouds is likely to be unbalanced (e.g. denser 3D points for objects close to the laser scanner, **Fig. 1d**), we improved a progressive triangulated irregular network (TIN) algorithm (Zhao et al., 2016) to interpolate the unbalanced point clouds. Then, we utilised all the above-ground points to generate a digital surface model (DSM), followed by the conversion of geo-coordinates on the × and y axes into pixel coordinates (Ritter and Ruth, 1997), defining four red-coloured ROI markers in the DSM (**Fig. 2c**). When processing a series of point cloud files collected from the same field, these four markers could be used repeatedly. To reduce computational complexity, we associated z values of each 3D point with a grayscale value (i.e. 0 cm is taken to be black, and 160 cm is taken to be white; the taller the point, the higher the grayscale value), followed by a projection method to cast all 3D points onto the flattened ground plane with an exchange rate of 1 cm per pixel. This process produced a 2D CHM image from an overhead perspective (**Fig. 2c**). Finally, we performed a 2D perspective transform (Mezirow, 1978) using the `getPerspectiveTransform` function in OpenCV (Howse, 2013) to extract the region within the four markers and align CHM images for automated trait analysis. The 2D CHM image contains spatial information for the experimental field and all the plots in the field.
4. *Steps 4*: to segment plots using the 2D CHM, we employed the 2D Hough transform (Duda and Hart, 1972) to detect plot boundaries. Because the gap between plots could be unclear during the season (e.g. lodging could cover the gap), missing pixels between plots or noise could affect the result of the Hough transform. Hence, we designed an improved method to detect horizontal and vertical lines separately (**Fig. 2d**), including: (1) combining both global (Sauvola and Pietikäinen, 2000) and local thresholding (Firdousi and Parveen, 2014) methods to establish an initial plot mask for the 2D CHM, even if the background is not uniform; (2) differentiating ground (i.e. soils) and non-ground (e.g. plots) pixels through the mask; (3) using the *Sobel* operator (Kroon, 2009) to detect the horizontal and vertical edges (angles were set at 360 and 30 as all the CHMs were aligned); (4) drawing straight lines based on the detected edges (with right angles and x- and y-intercept as input parameters) using the `hough_line` and `line_aa` functions in Scikit-Image (van der Walt et al., 2014); (5) merging multiple detected lines if they were close to each other, so that only a single line could represent the gap between plots (**Fig. 2e**); (6) finally, assembling the lines and producing a final plot-level mask to present all of the plots in the field (e.g. 486 plots in the 2019/2020 season). To remove edge effects, gaps within plots due to plant sampling, and crop variation that is not directly linked to the varieties or treatments (e.g. N loss), we calculated the weighted centroid of each plot using entropy features (Susan and Hanmandlu, 2013) based on the distribution of grayscale values (i.e. crop height values) in the plot. Through this approach, width and length of a plot mask could be adjusted adaptively to rectify the sampling areas (**Fig. 2f**).
5. *Steps 5&6*: the last two steps of the pipeline measured and exported key performance- and yield-related traits for each plot. A range of traits have been measured, including crop height, 3D canopy uniformity, 3D canopy surface, canopy coverage and biomass estimation (i.e. 3DVI and 3DPI). A table (in CSV format) was generated and populated with these scores, with each row corresponding to a plot (i.e. a variety) and each column corresponding to a trait, arranged according to the plot location (i.e. row and column IDs) in the field (**Fig. 2f**).

### The GUI of CropQuant-3D

To facilitate non-expert users to process 3D point clouds (in LAS format), we developed the GUI of CropQuant-3D, which integrated the above analysis pipeline into a single dialogue panel, from which all analysis results could be downloaded. The GUI was implemented using PyQt5, a comprehensive set of Python bindings for the Qt v5 library (Summerfield, 2015), allowing the GUI to be executable on varied operating systems (see **Availability and Requirements**). Following similar systems designs described previously (Zhou et al., 2017b; Zhou et al., 2017a), CropQuant-3D uses a stepwise approach to process point clouds and analyse 3D traits. The initial window (**Fig. 3a**) shows several sections with default input parameters pre-populated. In the input section, a user needs to select a LiDAR file (test LAS files provided on the GitHub). Then, the user needs to pre-process the selected file, including denoising and ground-based filtering (*Steps 1 & 2* in the GUI). After the pre-processing phase, the user can generate a 2D CHM (*Step 3*) by defining the exchange rate between a pixel (i.e. a 3D point) and a metric unit (i.e. cm), followed by defining geo-coordinates of the experimental field (i.e. ROI markers of the field; *Step 4*). The *Step 5* is to segment plots using the 2D CHM, so that traits such as height and canopy coverage can be measured (*Step 6*). Finally, if the user needs to export point clouds for specific plots, the user can click four corners of one or multiple plots in the CHM following the order upper-left, upper-right, lower-left and lower-right (Step 7). To enable a fast selection of plot-level 3D points, we used the `EVENT_LBUTTONDOWN` function in OpenCV to create a mouse response event. The analysis results can be downloaded after all the steps are accomplished (**Fig. 3b**).

**Figure 3:**
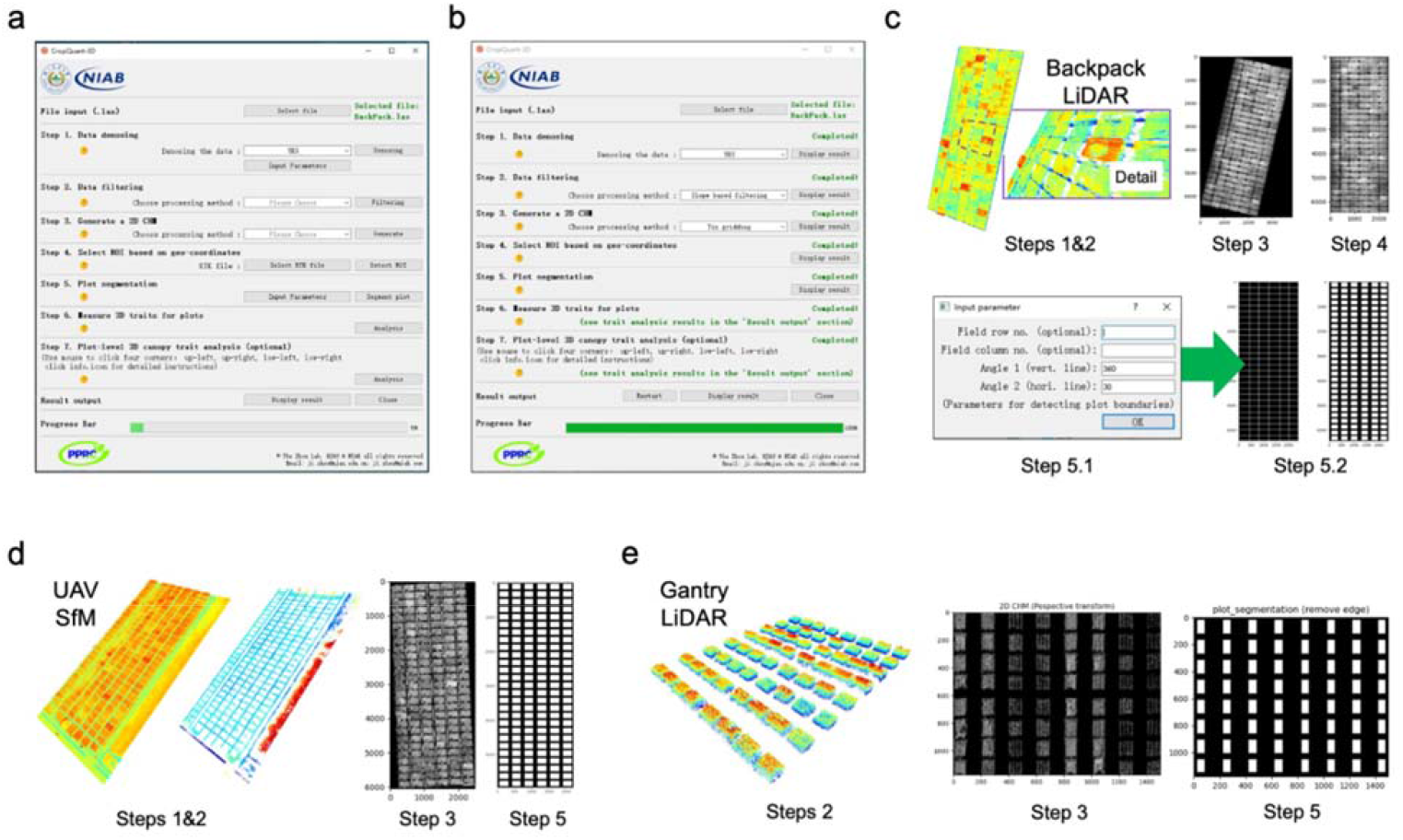
The graphical user interface (GUI) for CropQuant-3D designed for processing 3D point cloud files using 2D/3D image analysis algorithms and mathematic transformation for analysing canopy structural traits in 3D. (**a**) The initial GUI window of CropQuant-3D. (**b**) The GUI window after accomplishing all required analysis steps, with the progress bar showing 100%. (**c**) The intermediate results that can be displayed for each processing step integrated in the analysis procedure for processing point cloud files generated by the backpack LiDAR, including optional input parameters such as the number of rows and columns of the experimental field that users could enter to assist the algorithm for segmenting plots. (**d**) The intermediate results that can be displayed for processing point cloud files collected by UAV aerial imaging. (**e**) The intermediate results that can be displayed for processing point cloud files generated by gantry-mounted LiDAR systems.

When a step is finished, a green-coloured message will be displayed in the section together with a Display button to show intermediate results (**Fig. 3c**). In particular, if the gaps between plots are unclear and the plot segmentation (*Step 5*) fails to identify some plot boundaries, the user can define the field layout (i.e. the number of rows and columns) using optional input parameters, through which base lines will be generated to assist the segmentation algorithm. Furthermore, to enable the GUI software to process point clouds produced from other sources such as UAV-SfM and LiDAR mounted on gantry systems, we expanded the input function to accept these types of point cloud files (in LAS format). For example, the CropQuant-3D GUI can process point clouds generated by UAV-based aerial imaging (**Fig. 3d**) and FieldScan™ (Phenospex, Netherlands; **Fig. 3e**) following the same analysis steps integrated in the software to perform plot-based phenotypic analysis. A detailed step-by-step user guide (**Supplemental Materials S1**) and a video tutorial (**Supplemental Movie**) for the GUI software are provided in **Supplemental Materials**. The software implementation of the algorithmic steps is included in the **Materials and Methods** section.

### Height measurement using CropQuant-3D

Plant height and the rate of height increase (i.e. growth rate) are important performance- and yield-related traits (Holman et al., 2016; Nguyen and Kant, 2018; Momen et al., 2019). To measure crop height in a given plot, our algorithm was partially based on a mobile laser scanning approach described previously (Friedli et al., 2016), but performed on a flattened ground plane with the highest 10% 3D points (H_10_) in a given plot to reduce the height variances. The average height value of the H_10_ set was computed as the plot-level crop height. We produced three sets of height maps for all the 486 plots under three N treatments with a unified height scale bar (**Fig. 4**). The 3D DSM and 2D CHM images (**Fig. 4a-c**, left) show the 3D reconstruction and height distribution of the three N blocks, from 60-degree and overhead perspectives; whereas the coloured height maps (**Fig. 4a-c**, right) demonstrate how height of plants responded to different levels of N treatments (**Supplemental Materials S2**).

**Figure 4:**
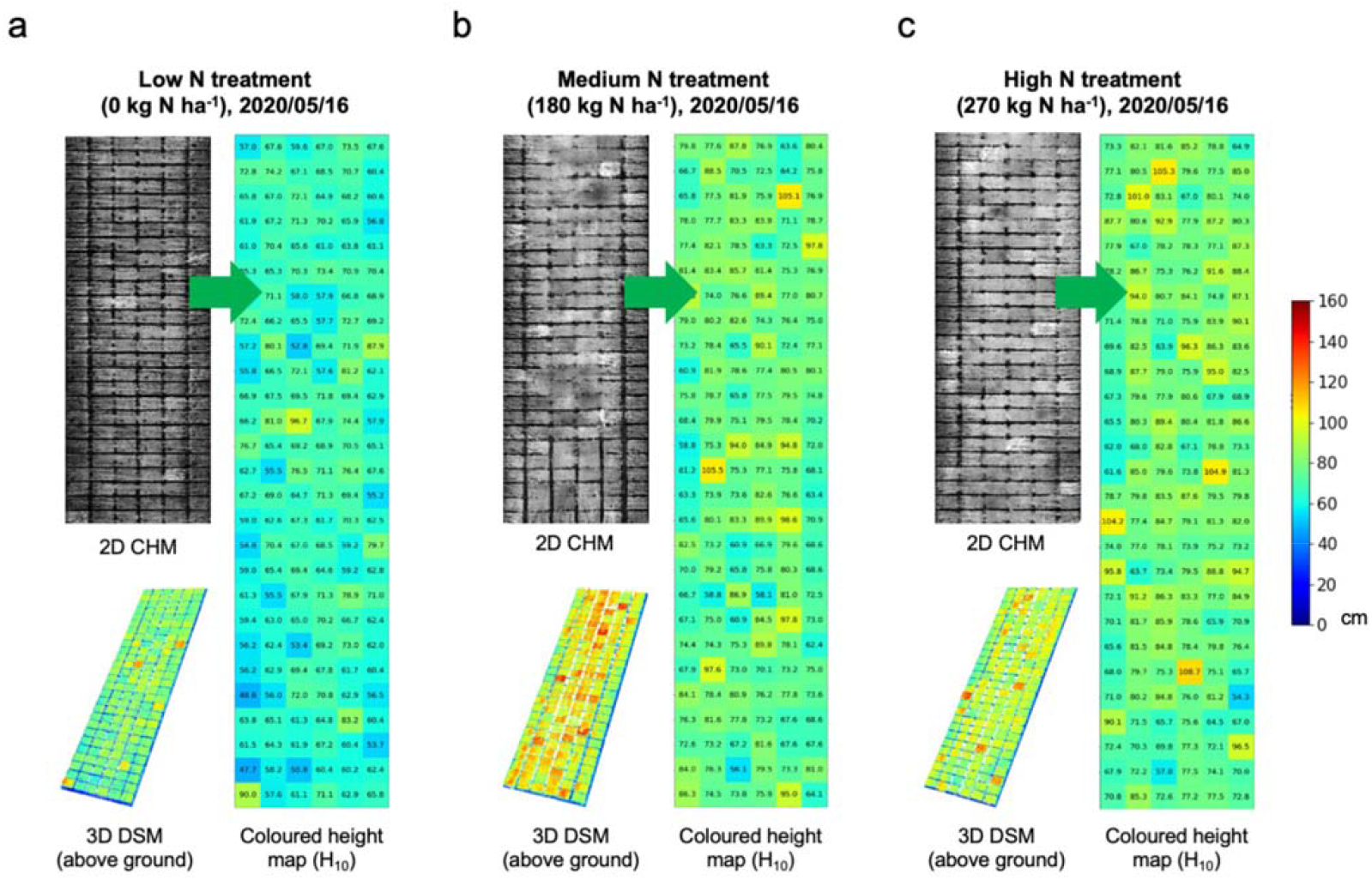
The pseudo-coloured uncalibrated height maps, 3D visualisation, and pseudo-coloured calibrated height maps of NUE wheat experiments under three different levels of N treatments. (**a**) The 2D CHM image (to the left) and 3D DSM image, created using the GPS-tagged altitude height values, and the calibrated height maps (to the right), showing the average height value of the highest 10% 3D points (H_10_) for the low N treatment; (**b & c**) the 2D CHM, 3D DSM (left) and the calibrated height (right) images for the medium N and high N treatments. The unified height scale bar for the three sub-figures is shown.

### 3D Canopy surface and 2D coverage measures

The rates of carbon gain through photosynthesis and water loss through transpiration of the canopy can be affected by changes in canopy structure, which can be used to explain crop performance and plants’ responses to environment (Green et al., 1985; Shearman et al., 2005). However, it is challenging to measure 3D canopy structure due to its complexity and dynamic spatial variability caused by genetic, agronomic management, and environmental effects (Omasa et al., 2007; Hosoi and Omasa, 2009; Duan et al., 2016). Although LiDAR devices have been used to visualise 3D canopy structure, we addressed the challenge of how to quantify canopy structural changes using point clouds. We initially approached the matter through measuring 3D canopy surface area and 2D canopy coverage traits using spatial features at the canopy level. To measure 2D canopy coverage index, we developed the following steps: (1) retaining highest 50% 3D points (H_50_) in a given plot (**Fig. 5a**); (2) then, projecting H_50_ points onto a flattened plane to generate a 2D canopy image from an overhead perspective; (3) after that, applying the `threshold_local` function in Scikit-Image (Singh et al., 2012) to select pixels in the canopy image using the calculated local threshold, resulting in a binarized canopy mask to represent the plot-level 2D canopy coverage. We applied the trait to measure canopy coverage differences of a wheat variety (e.g. NMzi-1019) under three N treatments. The index (0-1, where 1 is 100% coverage) showed an increase of around 10-15% when the N fertilisation increased (**Fig. 5b)**.

**Figure 5:**
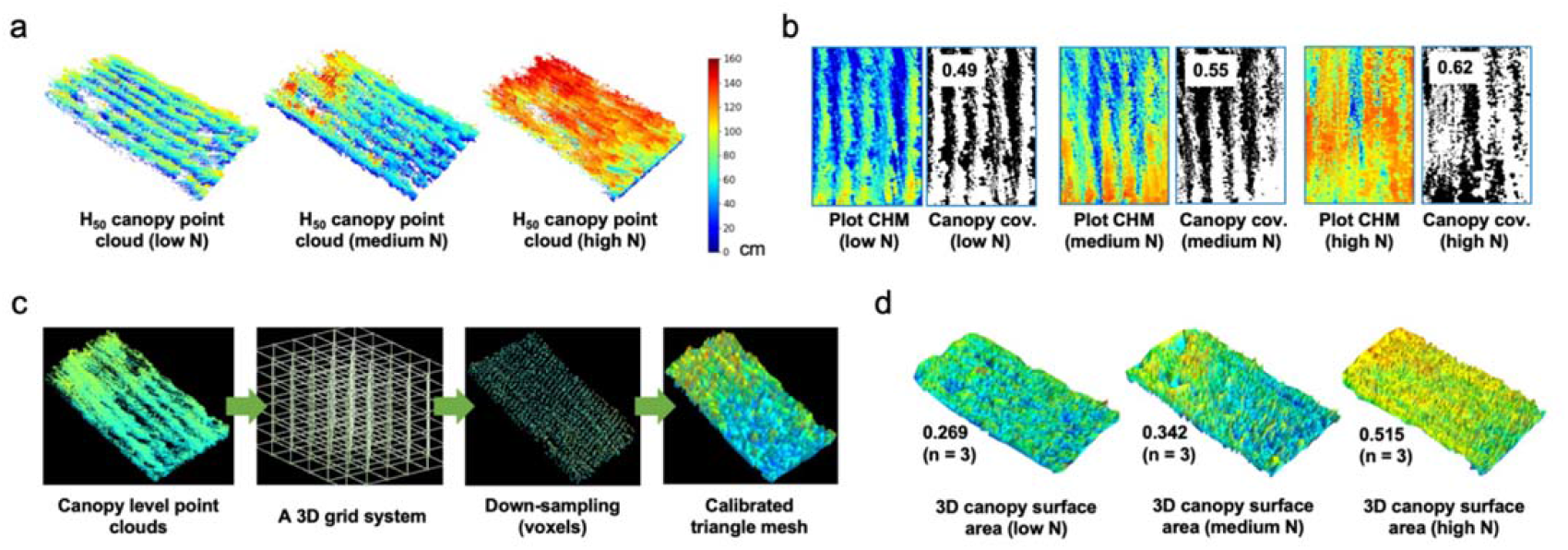
The analysis process of measuring 3D canopy surface area and canopy coverage at the plot level using voxels and triangular mesh for wheat varieties. (**a**) 3D points for the canopy region using the highest 50% points (H_50_) in a given plot. (**b**) H_50_ points projected onto the ground plane, generating pixels representing crop canopy regions, which were processed by an adaptive approach to calculate the canopy coverage trait. (**c**) The analysis process of computing the 3D surface area trait using triangle mesh. (**d**) The 3D surface reconstruction results of a wheat variety under three N treatments.

While the 2D canopy coverage area is important as it relates to the interception of direct solar radiation, it does not account for the total leaf area of the canopy, which is a more precise measure of interception of diffuse radiation and reflected light within the canopy (Cabrera-Bosquet et al., 2016). Also, the 3D surface area of the canopy would be closely related to the total transpirational leaf area than the 2D canopy cover, and would be more correlated with the summed photosynthetic activity of all leaves in the canopy region (Omasa et al., 2007). Therefore, we included 3D canopy surface area in the CropQuant-3D (**Fig. 5c**). The algorithmic steps were designed based on the triangle mesh method (Edelsbrunner et al., 1983), including: (1) applying the voxelization method (Truong-Hong et al., 2013) to generate a 3D grid system to package 3D points into voxels; (2), using the `voxel_down_sample` function from Open3D to down-sample the number of voxels, so that different parts of the plot (e.g. gaps between plants) could be covered; (3) using the `create_from_point_cloud_alpha_shape` function (Edelsbrunner et al., 1983) to reconstruct 3D surfaces of the crop canopy using the triangle mesh approach, followed by the `get_surface_area` function to calculate the 3D surface area. For example, the 3D surface area indices of wheat variety NMzi-109 showed an increase of over 20% with the increase in N application levels (**Fig. 6d)**. In addition to the above two traits, we also integrated traits such as 3D voxel index (3DVI) and 3D profile index (3DPI) into CropQuant-3D to estimate biomass, which has been described previously (Richardson et al., 2009; Jimenez-Berni et al., 2018; Deery et al., 2020). The results of 3D trait analysis are listed in **Supplemental Materials S3**.

**Figure 6:**
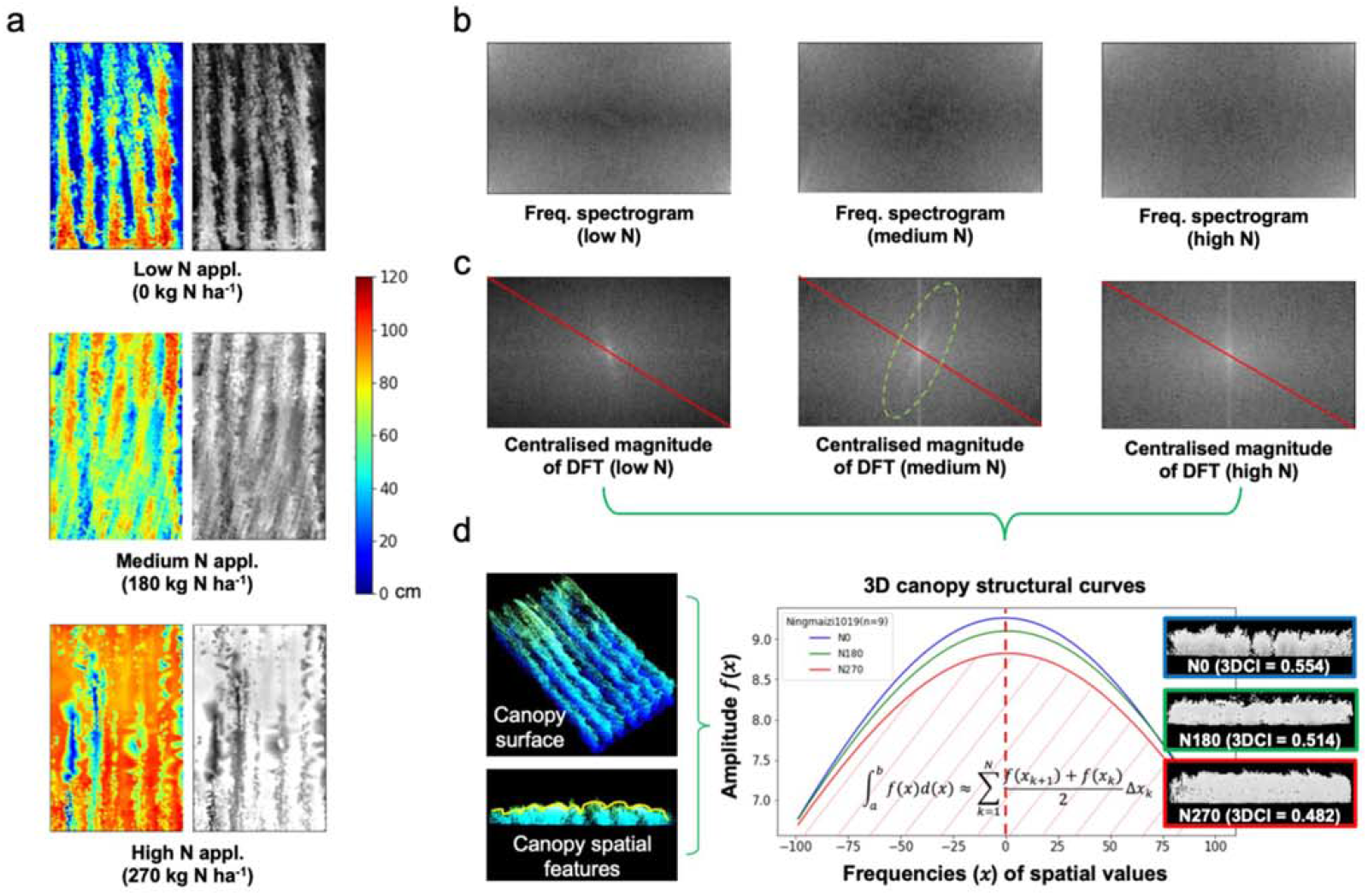
The analysis procedure of measuring 3D canopy structure at the plot level using 2D CHM images and a 2D discrete Fourier transform, resulting in 3D canopy structural curves for separating variety responses to different N treatments. (**a**) The pseudo-coloured height images and their associated grayscale height images (intensity values correspond to height values) in a plot, under three N treatments. (**b**) Frequency spectrograms generated using 2D Discrete Fourier Transform (DFT) of the grayscale height images, containing all frequencies of height values and their magnitude in the plot. (**c**) Centralised magnitude of DFT produced to enable frequency and amplitude sampling through red coloured lines on the diagonal of the image; regular patterns observable in the images with medium- and high-N treatments. (**d**) Three canopy structural curves plotted to present structural differences together with cross-sections of 3D points at the canopy level, showing the wheat variety’s different responses to three N treatments as well as the procedure of computing 3D canopy index (3DCI) based on the curves and areas beneath the curves.

### A new canopy structural measure – 3D canopy index

Whilst the above indices are useful measures to describe canopy structural features, they do not convey information about changes in spatial features (e.g. height) across the plot, which are likely to be affected by many factors in the field, including (1) individual tillers (e.g. main stem is taller than secondary tillers), which could differ between genotypes; (2) the height of spikes if a mixed population was drilled; (3) the density of the crop (e.g. spikes number per unit area, SN m^-2^) due to different management practices such as the seeding rate, (4) agronomic or environmental reasons unrelated to treatment or genotype (e.g. local seedbed variations), and (5) plant lodging due to tall plants. We have established a new algorithm incorporated in the CropQuant-3D software to measure height changes at the canopy level. Following the previous naming convention (Jimenez-Berni et al., 2018), we called this measure 3D canopy index (3DCI). The algorithm for 3DCI consists of five key steps:

1. Using the plot-level masks (**Fig. 2e**), we extracted all the above-ground 3D points in a given plot to generate a pseudo-colour spatial map from an overhead view. We then transformed the map into a grayscale image with each pixel’s grayscale value corresponding to its height value, resulting in a 2D plot-level CHM (**Fig. 6a**, right).
2. A 2D discrete Fourier Transform (DFT) method (Cooley and Tukey, 1965) was applied to represent the plot-level CHM in the frequency domain, producing the magnitude of the image’s Fourier transform. Because the dynamic range of the Fourier coefficients was too large to be visualised, we applied a logarithmic transform and generated a frequency spectrogram (**Fig. 6b**), containing all frequencies of the spatial information in the plot and their magnitude. The DFT can be defined as: 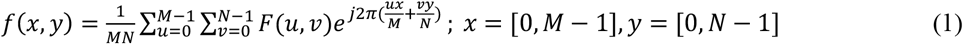 Where *f(x, y)* represents the M×N spatial domain matrix, and *F(u, v)* represents the DFT of *f(x, y)*. The coordinate system of *F(u, v)* is in the frequency domain.
3. We centralised the frequency spectrogram to remove periodic interference signals, resulting in a centralised magnitude image to represent the spatial information. For example, by applying DFT to CHM images under three N treatments, we could identify different structural features at the canopy level (**Fig. 6c**): (a) the magnitude of the low-N magnitude image became rapidly smaller for higher grayscale values (i.e. canopy objects such as spikes with taller height), suggesting its canopy was lower and the distribution of its spatial features was spread out (i.e. less dense) compared with crops under medium or high N treatments; (b) the main values of spectrogram images for both medium and high N applications lay on a vertical line, suggesting their canopy structures contained a dominating vertical orientation caused by regular spatial patterns (e.g. lines of plants); and (c) in the medium-N magnitude image, another pattern could be observed which passed through the center at 75-80° angle (highlighted by a light-green dashed oval), which was caused by another pattern in the plot and potentially could be a useful tool to measure the degree of lodging (**Fig. 6a**).
4. To utilise the above DFT results in quantitative trait measurements, we sampled all the pixels’ grayscale values on the diagonal of the centralised magnitude image (red coloured lines in **Fig. 6c**), based on which frequencies of all spatial values and their amplitude were summarised. We then used the Gaussian fitting (Virtanen et al., 2020) to plot the amplitude of the sampled spatial values, producing curves to represent canopy structural features within a defined frequency region (i.e. -100 to 100), where the x-axis denotes frequencies of canopy-level spatial values and the y-axis represents their associated amplitude (**Fig. 6d**). Two important features could be concluded from a given curve: (a) the curvature, signifying the density of crop canopy, as a less dense canopy structure resulted in a higher curvature due to larger spatial variations in the canopy; (b) the area beneath the curve (e.g. with red diagonal stripes, **Fig. 6d**), showing the canopy uniformity – when curvatures are similar, a curve with greater area indicates less uniformity due to greater accumulated spatial variances and hence a smaller density. To combine both curvature and area, we used integral calculus (i.e. integration) to compute the canopy curve area, which is defined by Eq.2: 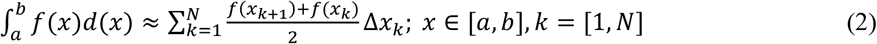 Where *x* is frequencies of spatial values, *a* is the minimum frequencies of spatial values (set as -100), *b* is the maximum frequencies (set as 100), *f*(*x*) is the amplitude value after Gaussian fitting, indicating the canopy uniformity, *N* is the total number of × sampled, Δ*x*_k_ is the difference between *x*_k_ and *x*_k+1_. To compute the curvature (Van Der Walt et al., 2011), we used Eq.3: 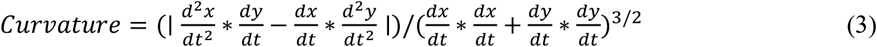 Where *x* represents the frequency array (the x-axis), *y* is the amplitude array (the y-axis) of the canopy structural curve.
5. To use the new metric for measuring canopy uniformity in CropQuant-3D, we normalised values generated by Eq2., so that we could cross-validate the measure for different plots. We called this normalised value 3DCI. The normalisation is defined by Eq4.:

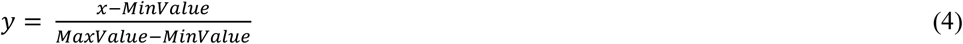

Where *x* is the calculated value, *y* is the normalised 3D canopy uniformity index, *MinValue* is the theoretical minimum value from the value list, i.e. 59.3% of the calculated minimum value (Raybould and Quemada, 2010); and *MaxValue* is the theoretical maximum value from the value list, i.e. 129.4% of the calculated maximum value.

To verify the 3DCI measure, we used the wheat variety NMzi-1019, which has shown to respond strongly to different levels of N fertilisation (Feng et al., 2008). Three canopy structural curves of NMzi-1019 for three N treatments (n = 3 plots) were produced (**Fig. 6d**). The three curves’ curvatures reduced moderately when the N fertilisation increased, indicating the canopy density were increasing. The high-N curve (coloured red) contained less spatial variations than those with low- and medium-N treatments (see cross sections in **Fig. 6d**) and possessed a smaller area beneath the red canopy curve. Trends in 3DCI scores across N treatments were used to differentiate varietal differences in canopy responses to N treatments. Increasing 3DCI indicated that the canopy became more variable in height and hence more structural changes at the canopy level. Similarly, if the index decreased with increasing levels of N, this indicates that the canopy became more uniform and likely much denser.

### Validation of the traits using ground truth data

Height estimates derived from the CropQuant-3D output were validated by comparisons with manual height measurements taken at the same stage of crop development (early grain filling) in the 2019/20 trial. There was a strong correlation between the CropQuant-3D’s height scores and manual field measurements for each level of N, using both plot-based (R^2^ ranges from 0.69 and 0.87, p < 0.001; **Fig. 7a**) and variety-based means (R^2^ ranges from 0.84 and 0.92, p < 0.05; **Fig. 7b**; **Supplemental Materials S4**). Thus, the CropQuant-3D height scores based on the backpack LiDAR provides a viable alternative to manual height measurements, particularly for obtaining genotypic means. It is also interesting that CropQuant-3D tended to underestimate the height for wheat varieties that are taller than 90 cm (some landraces were included in the field experiments). This is likely due to the way manual measurements were taken, which involved lifting and straightening curved or lodged plants to measure the distance from the soil surface to the tip of the ear along the vertical stem, whereas the LiDAR system measured the plants as they were naturally in the field. Furthermore, because only a limited number of plants were measured in each plot manually, compared with a whole plot scan conducted with the LiDAR system, there is greater chance of plot-to-plot variability with the manual approach than with LiDAR, which integrates height measurements over a larger number of plants in a plot. Also, better variety-based correlation values might be due to height values for each variety have been averaged (three replicates per variety), reducing the height variance caused by treatments and small agronomic differences.

**Figure 7:**
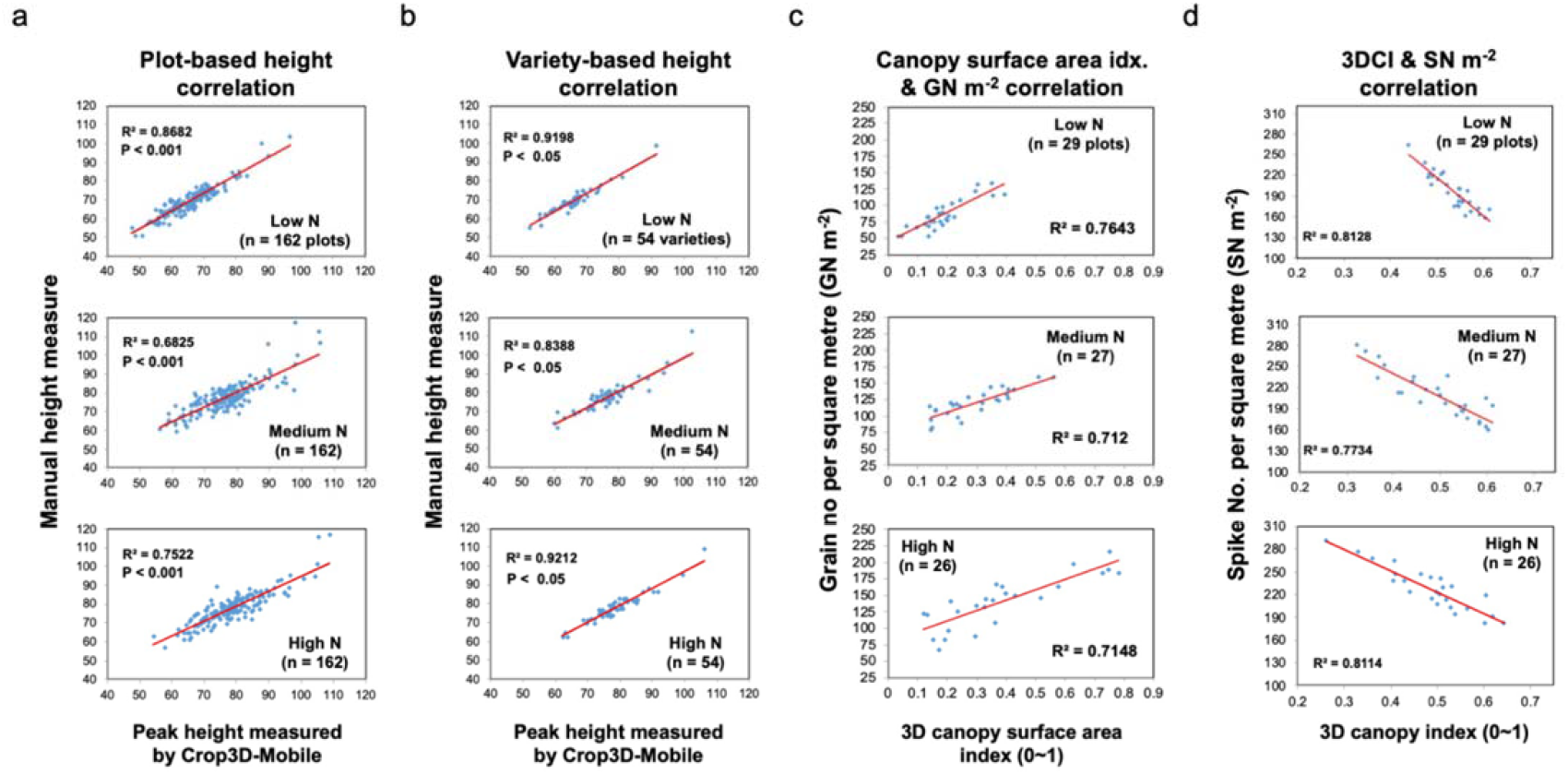
Correlations between height estimates, canopy surface index, and 3DCI computed by CropQuant-3D and manual measurements in the 2019/20 field trial, at three different levels of N fertilisation. Plot means (a) and genotype means (b-d) are shown. (**a**) Plot-based correlation analysis of the peak height measured by CropQuant-3D and manual height measurements. (**b**) Variety-based correlation analysis of the peak height measured by CropQuant-3D and manual height measurements. (**c**) Correlation analysis of the 3D surface area index and the grain number per unit area (GN m^-2^) data. (**d**) Correlation analysis between the 3D canopy index (3DCI) and spike numbers per square metre (SN m^-2^).

To verify the biological relevance of the 3D canopy surface area index, we have analysed correlations with plot-level grain number (GN m^-2^) and grain yield (GW m^-2^) using data from the 11 selected varieties (n = 81 plots). Strong positive correlations between this LiDAR-derived trait and the yield components, with R^2^ ranging from 0.71 to 0.76 (p < 0.001, **Fig. 7c**), suggest a mechanistic link between the 3D canopy trait and grain formation underlying the correlation, indicating that the 3D surface area index can serve as a good predictor of varietal performance. Additionally, there was a strong negative correlation between 3DCI (designed to quantify canopy uniformity) and manual measurements of spike density (SN m^-2^), R^2^ ranging from 0.77 to 0.81 (**Fig. 7d**; **Supplemental Materials S4**). Hence, it is highly likely that the 3DCI introduced here can also be utilised on a large scale as a measure to quantify how an important yield component responds to different N applications, but without the slow and laborious process of manually counting spikes in the field.

### A case study of classifying nitrogen responses for wheat

To effectively select crop varieties with an improved N response (e.g. high nitrogen use efficiency, NUE), it would be valuable to make use of proxy traits that are related to NUE under field conditions (Sylvester-Bradley and Kindred, 2009; Pask et al., 2012; Nguyen and Kant, 2018). The variables describing canopy structural features that measured by CropQuant-3D (i.e. 3D canopy surface area, 2D canopy coverage, plot height and 3DCI) were combined to enable us to classify the N response of 11 selected wheat varieties (81 plots) into four classes of N response pattern (**Fig. 8, Supplemental Material S5**). The example varieties were as follows:

**Figure 8:**
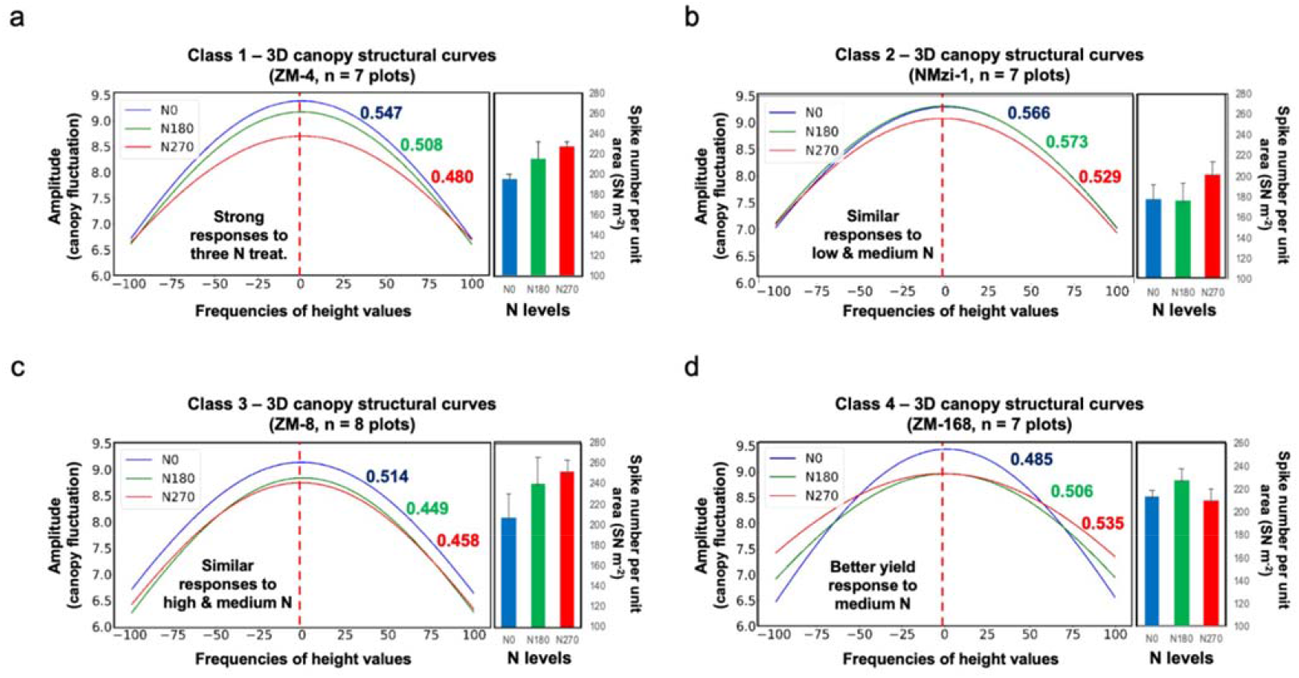
A case study of classifying wheat varieties’ nitrogen responses using the 3D canopy index and spike number per unit area for 11 varieties from the Zhenmai and Ningmai collections under three N application levels. Values shown in corresponding colour next to each curve in the plots are computed 3DCI values, which combines curvature and area under the canopy curve. (**a**) The first N response class, showing 3D canopy structural curves of ZM-4 and the associated spike number per metre square (SN m^-2^) scores under the three N treatments. Also in this class were varieties NMzi-1019, ZM-5 and ZM-1 (see text and Fig. 5 for the explanation of the measure). (**b**) The second N response class, showing 3D canopy structural curves of NMzi-1 and the associated SN m^-2^ scores under the three N treatments. Also in this class were NMzi-1, ZM-10 and ZM-12. (**c**) The third N response class, showing 3D canopy structural curves of NM-26 and the associated SN m^-2^ scores under the three N treatments. Also in this class was ZM-8. (**d**) The fourth N response class, showing 3D canopy structural curves of ZM-168 and the associated SN m^-2^ scores under the three N treatments. Also in this class was line ZM-09196.

1. Class 1 – canopy structural curves differed across all three N levels. The patterns for ZM-4 could be clearly separated under the three N treatments (**Fig. 8a**), indicating that this type of wheat variety had a strong response to varied N applications. Both 3DCI (coloured according to their associated N treatment) and the curvatures of the three canopy curves reduced steadily together with the increase of N, indicating that spike density and canopy uniformity were both rising in response to the escalation of N treatment. Also, the decrease of 3DCI also correlated with a continual increase of the SN m^-2^ reading. Other lines from the 11 varieties that can be categorised into Class 1 are NMzi-1019, ZM-5 and ZM-11.
2. Class 2 – canopy structural curves were similar at low and medium N levels, but differed at high N. The patterns for NMzi-1 showed that the line had a good response to increased N application, but only above the medium rate of N fertilisation. Both 3DCI and SN m^-2^ suggested that low and medium N had similar effects on the variety (**Fig. 8b**). The SN m^-2^ increased distinctly only under high N. Other lines that can be categorised into Class 2 are NMzi-1, ZM-10 and ZM-12.
3. Class 3 – canopy structural curves were similar at medium and high N levels. The patterns for NM-26 suggested that the variety had similar responses under medium and high N treatments, indicating the increasing N fertilisation was not able to increase the line’s spike density beyond the medium rate of N fertilisation (**Fig. 8c**). The other line that can also be categorised into Class 3 is ZM-8.
4. Class 4 – canopy structural curves decreased at high levels of N and showed the best response at medium N. Curvature patterns of ZM-168 indicated that the line had a similar canopy density at medium and high N treatments. The spike density was greatest at the medium N level, although the greatest 3DCI value was observed at the high N treatment (**Fig. 8d**). The other line that can be categorised into Class 4 is ZM-09196.

We combined the 3DCI, crop height, canopy surface index area with the yield components GN m^-2^ and SN m^-2^ to produce a performance matrix to understand crop responses to different N treatments in a unified manner. According to the matrix, each variety was ranked based on the performance of these measures and traits. For example, by calculating the deviation of them based on the trimmed mean values (i.e. 15% over the trimmed mean coloured dark red and placed in rank order 5; 7.5∼15% coloured red and placed in rank 4; -7.5∼7.5% coloured yellow and placed in rank 3; -15∼-7.5% coloured light green and placed in rank 2; and -15% below the trimmed mean coloured green and placed in rank 1), we could select lines with a desired performance under the three N treatments using a ranking system. In particular, for crop height, both very short and very tall were ranked undesirable (i.e. rank 1), whereas both GN m^-2^ and SN m^-2^ were given more weight (Langer and Liew, 1973) than other measures (*weight* = [0.25, 0.25, 0.2, 0.1, 0.2]). Through the ranking system, we concluded that: (1) for the low N treatment, ZM-168 demonstrated a more balanced score in terms of grain production and phenotype (**Fig. 9a**); for the medium N application, NM-26 ranked the highest (**Fig. 9b**); and, for the high N, NM-26 was scored the highest (**Fig. 9c**). Although this is only an initial attempt for selecting wheat varieties with desirable N responses using LiDAR-derived traits and key yield components, it is evident that the performance matrix could provide an objective approach to rank the varieties. Further validation and field studies using the above selection approach are ongoing and will be reported separately.

**Figure 9:**
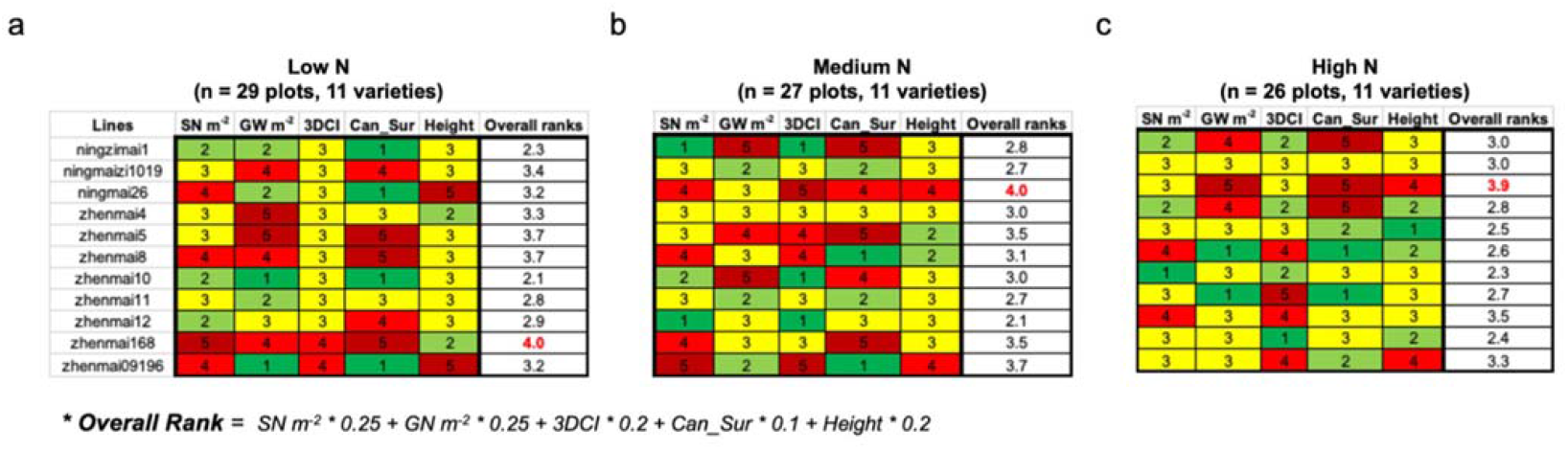
A performance matrix to evaluate NUE of wheat varieties using traits and measures for 11 wheat varieties from the Zhenmai and Ningmai collections under three N applications. (**a-c**) A range of canopy measures (i.e. 3DCI and canopy surface area index), plot level height, and key yield components (i.e. SN m^-2^ and GN m^-2^) combined to assess winter wheat varieties under three N treatments, with 15% over the trimmed mean coloured dark red, 7.5∼15% coloured red, -7.5∼7.5% coloured yellow, -15∼-7.5% coloured light green, and -15% below the trimmed mean coloured green. Selected varieties were coloured red, indicating they were ranked higher than the other varieties through the performance matrix.

## Discussion

Plant phenomics is an important area that helps provide valuable phenotypic information that is needed to fully exploit genomic resources. For crop improvement programmes, the focus is on multi-location field-based phenotyping approaches, yet there are a number of weaknesses with current solutions (Tardieu et al., 2017; Furbank et al., 2019; Pieruschka and Schurr, 2019), concerning: (1) mobility (a method can be straightforwardly used in multiple locations); (2) affordability (whether purchase, operation and maintenance of a system can be afforded by research groups with acceptable resources); (3) throughput (the number of plots, traits and fields that can be measured within a reasonable time frame, as well as the number of time points that can be acquired in a growing season); (4) accuracy (the information truly relates to the target attributes or biological functions of the plant); (5) resolution (if the method provides information at the level of detail quired to test the hypothesis); and (6) scalability (the size of trials that can be phenotyped and the number of locations that can be covered).

In addition to data collection, another issue that limits wide use of new field phenotyping tools involve the ability to analyse big and multi-dimensional data acquired from the field (Kelly et al., 2016; Scharr et al., 2016; Cendrero-Mateo et al., 2017; Lobet, 2017). Although many open-source and proprietary software solutions have been developed (Butler et al., 2020; Roussel et al., 2020), their applications are normally limited to certain devices and for questions that are mostly outside the plant research domain, leading to matters such as software usability, data interoperability, and the generalisability (Carpenter et al., 2012; Roitsch et al., 2019). To address some of the above issues, we pioneered the integration of backpack LiDAR for large-scale field phenotyping with GUI-based analysis software, CropQuant-3D, for measuring genotypic and N treatment differences in canopy structural features in wheat. Results from field experiments showed that canopy structural measures (e.g. height, 3DCI, and canopy surface area) are highly correlated with key yield components such as SN m^-2^ and GN m^-2^, indicating the system could be used as a reliable research tool to classify the plant responses to different N treatments.

### The backpack LiDAR hardware

We have shown that the backpack LiDAR device introduced here is integrated and portable, enabling the collection of high-density 3D point clouds at the field level. Typically, these kinds of data would require LiDAR systems to be mounted on a gantry or vehicle platform, which are often not available, too costly, fixed in one location, or cannot reach fields with limited accessibility. To our knowledge, the backpack LiDAR system has not been used in field-based plant phenotyping previously. Hence, we developed a range of techniques to apply the device for multi-location field. Our field testing and development experience show that the backpack LiDAR possesses three notable features: (1) large-scale capability (up to 210 m effective scan range through our equipment), with an acceptable mapping speed (up to 1.2 ha per hour); (2) portability (the ability to conduct multi-location phenotyping) with limited adjustments of hardware and software; (3) relatively small operation and maintenance costs due to its integration, ease-of-use and mobile features. Hence, backpack LiDAR appears to provide a more balanced solution to some current field phenotyping challenges. Although backpack LiDAR, like most high-resolution LiDAR systems with high-end scanners, is still relatively expensive. However, costs should decrease and become more affordable as the technology matures (Su et al., 2020). Comparisons between backpack LiDAR devices and other approaches can be seen in the section below.

### CropQuant-3D software and trait analysis

Processing of 3D point cloud data collected by LiDAR systems for trait analysis is still complicated and computationally demanding, indicating the necessity of reliable analytic solutions. Furthermore, for solutions that can be used by non-experts and widely accessible by the plant research community, the software should be user-friendly and open-source. Therefore, we developed the CropQuant-3D analysis software system to routinely process large point cloud datasets. To help other researchers exploit our analysis algorithms integrated in the software, besides the GUI software, we also modularised the analysis tasks into individual procedures and then saved them with executable Python source code in Jupyter notebooks that can be executed on multiple operating systems. The algorithmic steps include pre-processing of 3D point clouds (see code fragments in **Supplemental Materials S6**), automated plot segmentation with optional layout input, and plot-level crop height (see **Supplemental Materials S7**), 3D trait analysis of canopy structural features (3DCI, 2D canopy coverage, 3D canopy surface area), and biomass estimation (see **Supplemental Materials S8**). Compared with previously work (Ward et al., 2019; Hyyppä et al., 2020; Su et al., 2020), we have made progress in several areas for large-scale 3D trait analysis in plants:

1. Due to the huge volume of raw point cloud data collected, efficient data processing needs to be considered for both throughput and accuracy. Many existing methods require much computational time to pre-process point clouds. In our case, we have chosen to use a ground-level filter with parameters tailored for small-scale crop field, retaining only the 3D points required by trait analysis. This approach noticeably reduced processing time. For example, for a 400MB LiDAR file (over 15 million 3D points), only 100-120 seconds were required to normalise 3D points on an ordinary computer (intel i7 CPU and 16 MB memory; see profiling in the **Material and Methods**).
2. We analysed plot-level 3D traits using 2D CHM, which retains sufficient spatial information in 2D pixels. This approach enabled us to employ computationally efficient 2D-based algorithms such as edge detection, Hough transform, and adaptive thresholding to perform plot segmentation and trait analysis, reducing the computational complexity. Another key benefit for this 3D-to-2D transformation is that analysis regions could be controlled dynamically in the plot. By calculating the texture entropy (Haralick et al., 1973), we could compute the weighted centroid of a plot and then define the sampling area according to experimental needs.
3. Since the density of the LiDAR-collected 3D points is likely to be imbalanced (e.g. the further away from the mapping route, the sparser the 3D points), it is necessary to interpolate the point clouds if the number of 3D points in a given plot is limited. From a range of interpolation algorithms, we have chosen the progressive TIN to build a TIN-based model and then densify 3D points in an iterative manner, which helped us improve the quality of 3D trait analysis while retaining key 3D geometric features in the plots.
4. It is technically difficult to describe 3D canopy structure in a quantitative manner. The 2D Fourier transform method employed by CropQuant-3D opens a new door to quantify spatial variances and uniformity at the canopy level by dividing frequency and amplitude of all height values across the plot. A similar idea but with a different approach can be found in measuring the canopy roughness of leafy trees in forest ecology (Antonarakis et al., 2010). Our approach was able to show that, through the canopy structural curve and 3DCI (**Fig. 6d**), we could quantify the uniformity and density of wheat spikes in plots, which could be used to classify varieties according to different responses to N treatments. Meanwhile, the curvature of the canopy curves can also be employed to help distinguish the canopy density in relation to different N treatments and varieties (**Fig. 8d**).

There are many vision-based approaches developed to mine spatial and temporal features from point clouds for a range of biological questions, for example, identifying phenotypic differences at the organ level (Li et al., 2020a) and the extraction of single plants within a plot (Jin et al., 2021). Because our research aim was to enable large-scale field phenotyping for plot-level 3D trait analysis, we therefore did not consider plant-level 3D reconstruction and methods to analyse detailed features (e.g. plant-level marching cubes, leaf curvature estimation, and 3D skeletonization) in this work.

### Wheat varietal responses to different nitrogen fertilisation levels

NUE in crops is generally low. Approximately 40% of the applied N can be utilised by cereal crops, with the bulk of the remainder leaching to groundwater or volatilising to the atmosphere, causing increased agricultural costs and negative impacts on the environment (Raun and Johnson, 1999; Good et al., 2004). Breeding crop varieties with improved NUE should contribute to more sustainable cropping systems. To effectively select lines with heritable NUE-related proxy traits under different field conditions, efficient field phenotyping techniques are required (Nguyen and Kant, 2018; Nguyen et al., 2019). Nevertheless, it is technically difficult to screen many complex traits due to their dynamics and complexity (Good et al., 2004; Sylvester-Bradley and Kindred, 2009; Swarbreck et al., 2019).

In the case study, we have explored a comprehensive procedure to quantify N responses of different wheat varieties based on phenotypic traits and key yield components. When the level of N changed, different varieties varied with their responses in terms of canopy structural traits and yield components. By combining various yield component and LiDAR-derived trait values, we identified four NUE types using the subset of 11 varieties: (1) grain yield responded well to increased N application (Class 1); (2) only higher N was able to increase yield (Class 2); (3) medium and high N treatments led to similar grain production (Class 3); and (4) higher N led to a yield decrease (Class 4). We believe that the combined performance matrix demonstrated in the case study is likely to help establish an objective approach to identify wheat lines with superior N responses, which may lead to an effective selection improvement of NUE in wheat breeding programmes in the future. Further work to link this selection approach with yield production and NUE at a large scale is ongoing and will be reported separately.

### Applications of CropQuant-3D

The traits and measures here (e.g. height, coverage, canopy area, and 3DCI) do not just relate to N treatments, but they also closely connect with many aspects of genetic variation in crop performance. For example, crop height is an important factor in assessing risk to crop lodging, 3D canopy area and 2D ground coverage are good indicators for managing agricultural inputs to optimise canopy structure for radiation capture, photosynthetic output and transpirational water loss. It is also important to note that such canopy traits are only apparent in the context of a population in plots, and most of these traits are difficult or impossible to convey by phenotyping individual plants in controlled environments. Canopy traits are affected by variety, soil characteristics and agronomic factors such as seed spacing and the application of plant growth regulators. The accuracy of models that attempt to simulate the effects of these factors and their interactions on crop performance may be improved by supplying them with these kinds of canopy traits collected across a wide range of scenarios.

The 3D canopy traits derived from LiDAR data described here such as 3DCI have many underlying component traits and spatial features Better understanding of the bases of 3DCI would broaden its application for other crop improvement programmes. For instance, height variances within a plot could be due to a variety of reasons: 1) a mixed population of plants with different genes controlling height, or that major height genes are not fixed, but still segregating in the population; 2) agronomic or environmental variability within the plot that is not related to genotypes; and 3) as 3DCI is affected by height as well as spike density, it is likely that the analysis of 3D point clouds could pick up the differences in height of the mainstem and different tillers on each plant, and tillering responds both to N treatment and genotype (Power and Alessi, 1978). Another biological application of the CropQuant-3D system is for discovery of robust quantitative trait loci (QTL) for agronomic traits, which requires phenotypic data on large mapping populations across multiple field environments (Griffiths et al., 2012). The high-throughput capabilities of this system are well suited to this type of research. A similar approach has been reported in our recent work, SeedGerm (Colmer et al., 2020), which was applied to detect genetic differences in *Brassica napus* based on a range of germination traits. Although more work is needed, greater automation of phenotypic analysis and improvements in accuracy are likely to accelerate genetic analysis of crop performance under varied treatments or environments.

Beyond existing 3D trait analysis, continuous phenotypic analysis in 3D of different crop species is likely to extend our understandings of the physiological bases of crop growth and development, for which the open-source nature of CropQuant-3D is likely to be valuable for the research community. There is an additional analytic power in examining longitudinal traits (time-series measures of traits that change as the crop develops and matures), which can describe the dynamic nature of the interactions between crop genotype and N responses. By streamlining both the data acquisition and data analysis of field phenotyping with the backpack LiDAR and CropQuant-3D, it becomes possible to obtain measures at each key growth stage and at different test locations/environments, which was difficult to achieve with systems that are less portable and flexible in operation, with limited opportunity to expand or alter use of the analysis software. With this new approach, multi-environment 3D traits collected along a time series on large genotype collections can enable a deeper understanding of the genetic and physiological bases of efficient use of N for crop growth and development, and how these responses are modulated by the environment.

Technically, other than some supervised machine learning algorithms, we have not embedded popular deep learning techniques into the analysis pipeline for 3D traits analysis. Continuous development will improve our work, opening 3D phenotypic analysis to non-expert users and computational biologists that are willing to extend and jointly develop the system. Overall, we believe that the combined backpack LiDAR and CropQuant-3D system could have a great potential to advance large-scale and multi-location field phenotyping, 3D phenotypic analysis, and genetic studies for both crop research and breeding applications.

### Issues associated with the backpack LiDAR and CropQuant-3D

Despite certain clear advantages, it is also important to point out limitations of the backpack LiDAR and CropQuant-3D system. LiDAR technology has been maturing very rapidly in recent years. The Robin backpack LiDAR used in this study is already being replaced by new models with better accuracy, effective scan range, and a lower purchase price (the price of LiDAR devices has decreased over 30% since 2018; www.yole.fr/LiDAR_Market_Update_Livox_LiDAR.aspx). Although the backpack LiDAR is more affordable than other large-scale systems, it is worth noting that, depending on the laser scanner integrated in a backpack LiDAR device, the equipment is still relatively expensive. We compared the costs of Robin with some representative backpack LiDAR systems, as well as other LiDAR-based mapping approaches (**Table 1**; information regarding GPS and RTK accuracy can be found via the links in the References column). It is notable that the integration and mobility features of backpack LiDAR possess a unique opportunity for the community to explore shared services or community-driven facilitates encouraged by EMPHASIS and AnaEE (Roy et al., 2017).

**Table 1.** Cost comparison between backpack LiDAR devices, UAV airborne LiDAR, and the handheld laser scanning system, with brief technical specifications.

| LiDAR system | System costs<br>(academic price) | Brief technical spec. | References |
| --- | --- | --- | --- |
| Robin (backpack) | US\$350,000-375,000 with TerraSoild software (2018) | VUX-1 HA scanner, 1.5-200 m eff. range, 5-10 mm accuracy (outdoor) | www.3dlasermapping.com (discontinued) |
| BMS3D-HD (backpack) | US\$310,000-330,000 with BMS3D software (2021) | HDL-32 & VLP-16 scanners, 0.5-100 m eff. range, 20 mm accuracy (outdoor) | www.viametrisbusiness.com (France) |
| Pegasus (backpack) | US\$330,000-360,000 with local software (2021) | Dual VLP-16 scanner, 80 m eff. range, 10-30 mm accuracy (outdoor) | www.leica-geosystems.com (Switzerland) |
| GeoSLAM (backpack) | US\$300,000-320,000 with ORBIT software (2020) | Dual ZEB Discovery scanner, 100 m eff. range, 10-30 mm accuracy (outdoor) | geoslam.com (UK) |
| SLAM-based (backpack) | A low-cost solution with Forest3D software (2020) | Dual Velodyne Puck VLP-16 sensors, 100 m eff. range, 30 mm accuracy | (Su et al., 2020)<br>velodynelidar.com (US) |
| Airborne LiDAR | US\$125,000-150,000 with SpatialExplorer (2021) | SCOUT and RANGER series, 100 m range, 50-55 mm accuracy | www.phoenixlidar.com (US) |
| Handheld LiDAR | US\$25,000-50,000 with GeoSLAM Hub & Draw software (2021) | ZEB-Horizon scanner, 100 m eff. range, 10-30 mm accuracy | geoslam.com (UK) |

Additionally, our software was not designed to address many colour or spectral related traits that could be important for crop performance; for example, senescence of the lower canopy due to differential N or water limitation. Adjustments to how the LiDAR is used and the algorithms would be required to capture such traits. Similar issues can be applied to most of the LiDAR systems. It was difficult to scan the lower part of the crop after the canopy closure, which could cause errors to estimate above-ground biomass. We chose H_50_ and H_10_ for our canopy-level 3D analysis. Also, due to field conditions such as wind movement of the plants, it is extremely challenging to generate a very high-resolution 3D model to analyse an individual plant within the plot, through either increasing the accuracy of laser scanners or a close-up 3D mapping mode. Novel 3D points registration algorithms are needed to deal with plant movement to establish a reliable plant-level 3D modelling.

The CropQuant-3D system is capable of automating the segmentation of hundreds of plots for trait analysis, but the algorithm is likely to fail at the seedling development and tillering stages (GS10∼29). This is because the early crop height map and the gaps between drilled plants are too big to ensure a meaningful plot segmentation. However, as stems elongate and crop height increases (e.g. from the jointing stage onwards, GS31), our system can perform reliable plot-level masking. Another technical issue that needs to be taken into consideration is the request for a user to select plot(s) to extract plot-level point clouds. Although plot-level point clouds are not required for the trait analysis reported here, a user is required to select one or multiple plots on the 2D CHM to extract associated point clouds, which can be laborious if point clouds from hundreds of plots need to be extracted. For this technical constraint, automated plot-level 3D points extraction is required and recent reports suggest they are within reach (Walter et al., 2019; Roussel et al., 2020; Jin et al., 2021).

Because we have applied the 3D-to-2D analysis approach, some spatial information might be lost during the 3D-to-2D transformation, which could reduce the accuracy when the research interest is beneath the canopy region. For this loss of accuracy during 3D-to-2D transformation, we have performed some testing using 3D point cloud files collected by other equipment such as drone and vehicle mounted LiDAR (**Figs. 3d&e**). Although the preliminary is promising, further development and testing are still required to make the platform more compatible with these types of point cloud data. Next steps of the research also need to expand the application of CropQuant-3D to the analysis of different crop species so that the algorithms developed for wheat can be used for addressing similar biological problems in other crop species.

### Conclusion

Obtaining accurate and meaningful measures of the field phenotype at sufficient scale, throughput, cost and multiple locations create a bottleneck in today’s crop research and breeding, which is preventing us from making full use of genomic resources for crop improvement programmes. Backpack LiDAR has obvious advantages for large-scale field experiments and breeding trials. The device is easy to transport and use, overcoming the main limitations of a fixed phenotyping platform and can be used for multi-site data collection and at multiple time points. Another limitation of large-scale phenotyping is the ability to process and analyse large datasets with minimal time and standard computing power. To address this, we have developed CropQuant-3D, which processes large LiDAR-derived 3D point cloud data and consists of new algorithms packaged into a user-friendly GUI software to output multiple 3D canopy traits (e.g. 3DCI) at the plot level. In a case study of 11 wheat varieties grown under three levels of N inputs, Data obtained using the integrated backpack LiDAR and the CropQuant-3D software system showed that wheat varieties could be classified into different N response groups according to a range of 3D canopy structural traits that relate to crop height, spike density (SN m^-2^) and grain yield. This indicates that CropQuant-3D could be a useful tool to make selections for NUE, and to dissect the physiological mechanisms and genetic regulation of NUE. Hence, we trust that the combined backpack LiDAR and CropQuant-3D system has a great potential to relieve some of the current bottleneck in large-scale field phenotyping for crop research and breeding.

## Materials and Methods

### Plant material and field experiments

In the first season (2018/2019), 105 Chinese winter wheat varieties were planted at the Zhenjiang Agricultural Technology Innovation Center (ZATIC, 31°57’N, 119°18’E, Jiangsu province, China), measured using CropQuant-3D and assessed for yield and N responses. A subset of 54 varieties (**Supplemental Materials S9**) were chosen out of the 105 lines for the 2019/2020 season. The selected 54 Chinese winter wheat varieties used in the field experiments were cultivated from the wheat plantation regions of the middle and lower reaches of the Yangtze river, which were shown previously to vary in performance and yield under different nitrogen (N) treatments (Feng et al., 2008). A split-plot design was used, with three levels of N fertilisation as main plots, containing three replicates of the 54 varieties as sub-plots (162 plots per N experiment). The overall size of the 2019/20 field trial was 486 plots, covering approximately 0.5 ha (**Fig. 1a**). For the purposes of explaining the methods, data from 11 of the 54 varieties are shown.

### Crop management

Before sowing, soil samples (for 0-25 cm soil layer) were measured to ensure that available N content was suitable for N response studies (**Table 2**). Following standard crop management guidelines (Godwin et al., 2003) and local practice, base fertiliser (P_2_O_5_ and K_2_O) was applied before drilling. Three levels of N fertiliser treatments were applied by hand (0, 180, and 270 kg N ha^-1^) in two splits: 50% at sowing and 50% at jointing (GS31). Crops were planted in 6 m^2^ plots (2×3 m), with 6 rows per plot at 15 cm spacing, with approximately 30 cm gaps between plots (Fig. 1a; trial plans in **Supplemental Materials S9**). The planting density was 2.4 million plants per hectare. Plant growth regulator was not applied throughout the season so that stem elongation could respond unimpeded to different levels of N treatments.

**Table 2.** Soil nutrient (0-25 cm soil layer) content before drilling in the 2019/2020 season

| pH | Organic matter (g/kg) | NO<sub>3</sub>-N (mg/kg) | Phosphate (g/kg) | Potassium (g/kg) | Total organic N (mg/kg) | Plant available Phosphate (mg/kg) | Plant available Potassium (mg/kg) |
| --- | --- | --- | --- | --- | --- | --- | --- |
| 6 | 24.2 | 1.35 | 0.61 | 13.2 | 10.4 | 3.04 | 160 |

### Manual measurement

To collect reliable ground truth data for validating and improving CropQuant-3D’s analysis algorithm, a team of five field workers performed the manual scoring. They conducted a range of manual measures at key growth stages (from heading, GS51-59, to grain filling, GS71-89), including plant height, growth stage scoring, and key yield components such as spike number density (SN m^-2^), spikes per plant, grain number per unit area (GN m^-2^), and thousand grain weight (TGW). For example, manual plant height measures of five typical plants per plot were conducted on 11^th^, 18^th^ and 26^th^ May 2020, from which the scores on 18^th^ May (two days after the LiDAR mapping, 16^th^ May 2020) were used for correlation studies in this work. As there were variances in height across the plot, three one metre-square regions were selected to represent height variances within a plot. Then, all plants in the region were measured and the average height value was recorded as the plot height value. When measuring the plant height, the distance from the ground to the top of the ear was measured with a steel ruler. We took steps to standardise manual measurements: (1) cross-scoring same traits with different field workers, (2) cross-validating scores across experiments using historic data, and (3) using trimmed mean to remove outlier values before calculating the average of ground truth. At maturity, yield was measured in a 1 m^2^ quadrat centred in the plot, from which ears were removed with a sickle. Threshing was carried out with a plot thresher; any grain that passed through the thresher were manually recovered from the sieved straw.

### The backpack LiDAR system

The backpack LiDAR (Robin Precision, 3DLasermapping; purchased by GeoSLAM, Nottingham, UK) integrates a laser scanner (RIEGL VUX-1) and three mapping settings, employing accurate GPS-tagged navigation, and was used in conjunction with a real-time kinematic (RTK) base station for precise positioning. The system is a lightweight (around 10 kg) and comprises high-performance laser mapping system (360° scanning angle with an effective scan range of 3-200 m; further detail in **Supplemental Materials S10**). Measurements focussed on the key growth stages (Zadocks et al., 1974), from heading (GS51-59) to grain filling (GS71-89) when canopy structural features were largely established. Standard pre-processing software packages were bundled with the device. To capture the peak height for the selected wheat varieties, the trial was mapped from April to May 2020. In our preliminary work, similar 3D field mapping was conducted routinely in paddy rice trials at the Tuqiao crop breeding and cultivation centre (Jiangsu China) and at the Chinese Academy of Sciences’ (CAS) Songjiang crop research center (Shanghai China, **Supplementary Materials S11**). Notably, CropQuant-3D is not bundled with Robin and can be used to analyse point cloud files generated by other sources.

### GUI-based software development

To develop the GUI-based analysis software for CropQuant-3D, we utilised PyQt5, a comprehensive set of Python bindings for the Qt v5 library (pypi.org/project/PyQt5/), which was developed using C++ and is cross-platform for modern desktop (e.g. Windows and Mac OS) and mobile (e.g. Android and iOS) systems. The GUI software we developed follows a traditional desktop-based user interface development, which can be easily modified to operate in a web browser such as Google Chrome. Anaconda Python release (docs.continuum.io/anaconda/install/windows) was employed as our integrated development environment, through which third-party libraries required for the software implementation, testing and packaging were managed by multiple virtual environments installed into the `conda` directory (Virtanen et al., 2020). Algorithms (in Jupyter notebooks), GUI software (in EXE format), Python-based source code and testing files (in LAS format) are freely available.

### Software implementation

To implement *Step 1* (denoising) in the analysis pipeline introduced in the **Results** section, we first used the **file**.**File** function in the laspy library to read the input file, followed by the **spatial**.**cKDTree** function in the Scipy library to index the 3D coordinates of all the points in the LAS file. Then, we applied the filtering criteria (i.e. *avg*. + *k* × *std*.) to index outliers in the point clouds and saved the denoised point cloud data using the function **file**.**File** (in LAS format).

For the *Step 2* (filtering) in the pipeline, we developed three approaches to process point cloud files generated through different approaches: (1) for the backpack LiDAR mapping, we used the function `lidar_ground_point_filter` in the WhiteboxTools library to filter the point cloud; (2) for UAV-SfM generated pint cloud files, we employed the function `do_filtering` in the CSF library to separate ground-level 3D points from above-ground points; (3) for the gantry-mounted LiDAR files, because the 3D points have already been filtered, we could use the files directly.

For the *Step 3* (the generation of CHM) in the pipeline, we also developed three approaches to process different types of point cloud files: (1) for the backpack LiDAR generated files, we applied the function `lidar_tin_gridding` in the WhiteboxTools library to output CHMs with the resolution parameter set as 1 cm/pixel; (2) for UAV-SfM files, we used the `lidar_tin_gridding` function to output digital earth model (DEM) and DSM, followed by the `clip_raster_to_polygon` function to rectify the DSM and DEM’s resolution using the shapefile (the .shp file collected by RTK), resulting in an CHM imaging produced through subtracting the DEM from the DSM; (3) for the gantry LiDAR files, the `lidar_nearest_neighbour_gridding` function was used to produce the CHM image.

For the *Step 4* (the definition of ROI) in the pipeline, we used the function `read_csv` in the pandas library to read the geo-coordinates of the point cloud files, followed by the `open` function in the rasterio library to open the CHM and convert the geo-coordinates to pixel coordinates so that 3D point clouds could be analysed in 2D. The function `getPerspectiveTransform` in the OpenCV library was employed to obtain the perspective transformation matrix together with the `warpPerspective` function in OpenCV to define the ROI in the 2D CHM. Finally, the **io**.**imsave** in the scikit-image library was used to save the aligned 2D CHM within ROI.

For the *Step 5* (plot segmentation) in the pipeline, the optional input parameters such as the number of rows and columns could be used to generate horizonal and vertical base lines to assist the plot segmentation. Using the `threshold_sauvola` and `threshold_local` functions in scikit-image, we could obtain the threshold mask of the CHM image. Then, we applied the `sobel` function in scikit-image to detect edges in the CHM, followed by the `hough_line` function to fit vertical and horizontal lines, separately. By merging the detected lines and base lines, we could generate the final mask representing the plot boundaries in the field.

For plot-based 2D/3D trait analysis, we mainly used the `regionprops` function in scikit-image to calculate phenotypic traits in each plot. The plot-level 3D canopy traits were based on the `clip_lidar_to_polygon` function in WhiteboxTools to crop plot-level point clouds. The source code produced from the above software implementation can be seen in **Supplemental Materials S6-S8**, as well as from our GitHub repository.

### Software profiling

We profiled the GUI software using a range of testing point cloud files (in LAS format, available on our GitHub repository), which were acquired by the backpack LiDAR (403 MB; 15,090,552 points), UAV SfM generated point clouds (596 MB; 18,372,420 points), and gantry LiDAR (FieldScan™, 1.42 GB; 58,446,207 points). Three Windows laptop computers with different hardware configurations were used for the software profiling: (1) Intel Core i5 with 8GB memory (budget laptop); (2) Intel Core i7 processor and 24GB memory (middle-end laptop); and (3) Intel Core i9 with 32 GB memory (high-end laptop). As the CropQuant-3D software did not support GPU acceleration, both CPU and memory influenced the processing performance of CropQuant-3D. By averaging the computational time (using the `time` module in Python) used by the three computers, we provided details on the processing time using the three types of testing files at each step (**Table 3**).

**Table 3.** Processing time for three types of point cloud files at each analysis step

| Analysis steps | Processing time<br>(backpack) | Display time<br>(backpack) | Proc. time<br>(UAV) | Disp. time<br>(UAV) | Proc. time<br>(gantry) | Dis. time<br>(gantry) |
| --- | --- | --- | --- | --- | --- | --- |
| *Step 1* | 2.5-3 minutes | 35-45 seconds | 1.5-2 mins | 25-30 seconds | N/A | N/A |
| *Step 2* | 1.5-2 mins | 50-60 sec. | 30-40 sec. | 40-50 sec. | N/A | N/A |
| *Step 3* | 10-15 sec. | 0.2-0.5 sec. | 30-40 sec. | 0.5-1 sec. | 10-15 sec. | 0.3-0.5 sec. |
| *Step 4* | 0.5-1.5 sec. | 0.3-1 sec. | 0.5-1 sec. | 0.5-1 sec. | 0.5-1.5 sec. | 0.5-1 sec. |
| *Step 5* | 30-40 sec. | 0.5-1.5 sec. | 20-25 sec. | 0.5-1 sec. | 5-10 sec. | 0.5-1 sec. |
| *Step 6* | 5-6 sec. | N/A | 3-4 sec. | N/A | 1-2 sec. | N/A |
| *Step 7* | 5-10 sec. per plot | N/A | 4-6 sec./plot | N/A | N/A | N/A |

## Availability and requirements

Project name: 3D field phenotyping for wheat using backpack LiDAR and CropQuant-3D

Project home page: https://github.com/The-Zhou-Lab/LiDAR

Source code: https://github.com/The-Zhou-Lab/LiDAR/releases

GUI software: https://github.com/The-Zhou-Lab/LiDAR/releases

Programming language: Python 3.7

Requirements: Laspy (1.7.0), Whitebox (1.3.0), GDAL (3.1.4), Rasterio (1.1.8), Open3D (0.11.2), Mayavi (4.7.2), Scikit-Image (0.17.2), OpenCV-Python (4.4.0.46), Pandas (1.1.5), Numpy(1.19.4), Matplotlib(3.3.3), and Scipy (1.5.3).

License: BSD-3-Clause available at https://opensource.org/licenses/BSD-3-Clause

### Abbreviations

CSV: Comma-separated values
NUE: nitrogen use efficiency
IoT: internet of things
UAVs: unmanned aerial vehicles
LiDAR: Light Detection and Ranging
3D: three-dimensional
CT: computed tomography
RGB: red-green-blue
GNSS: global navigation satellite system
CHM: canopy height model
ZM: Zhenmai
NM: Ningmai
OS: operating system
N: nitrogen
SN/m^2^: spikes number per unit area
GN/m^2^: grain number per unit area
TGW: thousand grain weight
RTK: real-time kinematic
GPS: global positioning system
ROI: regions of interest
kNN: k-nearest neighbours
GUI: graphical user interface
RMSE: root-mean-square error
DFT: discrete Fourier transform
3DVI: 3D voxel index
3DPI: 3D profile index
R&D: research and development.

## Availability of supporting data

The datasets supporting the results presented here are available at https://github.com/The-Zhou-Lab/LiDAR/releases. Source code and other supporting data are also openly available on request.

## Author contributions

JZ and EO wrote the manuscript with inputs from YLZ, SCJ, QZ and JC. JZ, YLZ and GS designed the 3D phenotypic analysis algorithms. YLZ, GS and JieZ implemented the software under JZ’s supervision. GHD and MXW performed the field wheat experiments under JZ and YFD’s supervision. SCJ provided LiDAR expertise. JZ, YLZ and JieZ optimised the algorithms and tested the software. JZ, YLZ, QZ and JC performed the data analysis. All authors read and approved the final manuscript.

## Funding

The wheat field experiments are supported by the National Natural Science Foundation of China (32070400 to JZ). GHD and the field team were supported by Natural Science Foundation of Jiangsu Province (BK20191311 to JZ). JZ was partially funded by United Kingdom Research and Innovation’s (UKRI) Biotechnology and Biological Sciences Research Council (BBSRC) Designing Future Wheat Strategic Programme (BB/P016855/1) to Graham Moore. JieZ, SCJ and YFD were supported by Jiangsu Collaborative Innovation Center for Modern Crop Production. QZ was supported by Chinese Academy of Sciences (XDA24020205). JC was supported by the BBSRC’s National Productivity Investment Fund CASE Award, hosted at Norwich Research Park Biosciences Doctoral Training Partnership (BB/M011216/1). YLZ and GS were supported by the Fundamental Research Funds for the Central Universities in China (JCQY201902).

## Acknowledgements

The authors would like to thank all members of the Zhou laboratory at the Nanjing Agricultural University (NAU, China) and the Cambridge Crop Research, National Institute of Agricultural Botany (NIAB, UK) for fruitful discussions and cross-disciplinary collaborations. In particular, the authors would like to thank Prof Roger Sylvester-Bradley at ADAS, Dr Stéphanie Swarbreck at NIAB, and Prof Simon Griffiths at the John Innes Centre (JIC) for proofreading and helping us improve the manuscript. We also thank Prof Han Bin at the Chinese Academy of Sciences, Prof Dong Jiang, Prof Xiu-e Wang, Prof Daolong Dou, and Prof Yuqiang Liu at NAU for providing valuable suggestions for field experiments and data analysis. We thank researchers at the JIC and the University of Cambridge for constructive suggestions. We gratefully acknowledge the hardware support of the Zealquest Scientific Technology (Shanghai China) for the backpack LiDAR used in this research.

## Competing interests

The authors declare no competing financial interests.

